# Histone demethylase LSD1 regulates lipid homeostasis during *Cryptococcus neoformans* infection

**DOI:** 10.1101/2024.08.06.606889

**Authors:** Gaurav Kumar Lohia, Awantika Shah, Kithiganahalli Narayanaswamy Balaji

## Abstract

An opportunistic fungal pathogen, *Cryptococcus neoformans (C. neoformans)*, causes cryptococcal meningitis and is frequently associated with high mortality in immunocompromised individuals, particularly in HIV patients. Formation of metabolically altered lipid-rich foamy macrophages has been reported as a successful strategy employed by various intracellular pathogens to secure a nutrient source and niche within the host. Herein, we elucidate the involvement of macroautophagy, specifically lipophagy, in lipid dysregulation during *C. neoformans* infection. Our study highlights a pivotal role of lipophagy during infection, showing that *C. neoformans* driven activation of WNT-signaling leads to an aberrant lipid accumulation in host macrophages under the regulatory role of a histone modifier, Lysine Specific Demethylase 1 (LSD1). In a murine model of pulmonary infection, targeting host LSD1 led to a significant reduction in lung fungal burden, accompanied by amelioration of lung pathology and reduction of lipid content in the lungs. The study highlights the significance of host epigenetic regulation in modulating foamy macrophage formation through the regulation of lipophagy during *C. neoformans* pathogenesis.

## Introduction

*Cryptococcus neoformans (C. neoformans),* causative agent of cryptococcosis, accounts for 223,100 cases globally, with 181,100 deaths annually^1^. Recently, the incidence of infections caused by this fungus have increased at an alarming rate, correlating with a rise in the population of immunocompromised individuals^2^. Infection with *C. neoformans* primarily results in pulmonary associated pathogenesis, which under severe conditions (primarily immunocompromised conditions) disseminate to central nervous system wherein it manifests itself by causing fatal Cryptococcal meningitis^3^.

*C. neoformans* is reported to modulate various arms of host immunity to establish a conducive environment in the host. Among the various strategies opted, metabolic rewiring of host cells such as altered tricoboxylic acid (TCA) cycle, fatty acid metabolism, dysregulated glycolysis is garnering much attention in the recent times^4,5^. Augmented levels of lipids, in the form of Lipid Droplets (LDs), lead to the formation of metabolically altered, Foamy Macrophages (FMs) which have been implicated in various pathophysiological conditions. FMs exhibit impaired immune functions and aid in the disease pathogenesis by stimulating inflammation and tissue damage ^6^. To our interest, supplementation with exogenous oleic acid has been shown to enhance the intracellular growth of *C. neoformans*, suggesting a pro-pathogenic role for lipids^7^. Moreover, depletion of cholesterol, another major component of lipid droplets, has been reported to cause reduction in phagocytosis and clearance of *C. neoformans* in macrophages ^8^. Studies suggest *C. neoformans*-mediated induction of atherogenesis in the lungs of infected rabbits ^9^, as well as an increase in the uptake of lipid droplets by rat alveolar macrophages ^10^. The accumulation of triacylglycerol (TAG)-rich lipid droplets in macrophages has also been observed in the lungs of mice infected with *C. neoformans* ^11^.

In line with these observations and increased incident of *C. neoformans* infections, it is pertinent to understand the regulation of various cellular processes that could influence host lipid content during *C. neoformans* infection. In general, an increase in lipid content of the cell would be accounted by either increased biosynthesis and uptake of lipids or suppression of cellular processes involved in maintaining lipid degradation/ turnover ^12^. Lipophagy is the autophagic degradation of lipid droplets present in the cytoplasm for energy requirements during homeostasis or under stress conditions^13^. Defective or aberrant lipophagy promote the accumulation of lipid in the cytosol and has been implicated in many metabolic diseases such as fatty liver diseases, liver fibrosis and obesity^14^. Moreover, formation of lipid laden foam cells has been reported to occur via suppression of autophagic process^15^ and we have shown in our earlier studies that *Mycobacterium tuberculosis*, inhibits host lipophagy via modulating host epigenetic factors to promote foam cell formation and intracellular survival^16^.

Multiple studies evidence to infection induced epigenetic changes in the host cells via manipulation of key immune signalling pathways^17–19^. JMJD3, a Lysine demethylase, has been reported to play a crucial role in foam cell formation during mycobacterial pathogenesis^20^. H3K4 trimethylation was found to be necessary for human monocyte differentiation into foam cells^21^. In this perspective, Lysine Specific demethylase 1 or LSD1 (KDM1A) is known to support adipogenesis^22^, regulate lipid metabolism^23,24^ and is pivotal for efficient binding of Sterol regulatory binding protein-1 (SREBP-1) to its target gene promoters (Fatty acid synthesis genes), therefore, plays a role in activating lipogenic gene expression in mammalian cells^25^. Moreover, a role for LSD1 in viral infections, inflammation has also been underscored^26^. A flavin adenine dinucleotide (FAD)-dependent amine oxidase, LSD1, demethylates mono- or di-methylated histone H3 lysine4 (H3K4) and H3 Lysine 9 (H3K9) via a redox process leading to gene repression or activation respectively^27^. Therefore, our study focused on understanding the role of LSD1 in FMs generation during *C. neoformans* infection and its implications on pathogenesis.

Here, we report that *C. neoformans* infection results in augmented lipid content in macrophages *in vitro*. Mechanistically, we have observed a dominant role played by WNT-driven epigenetic modifier, LSD1, in transcriptional regulation of genes implicated in lipid uptake and biosynthesis. Furthermore, lipophagy was also found to be negatively regulated by LSD1 during *C. neoformans* infection. Utilizing an *in vivo* prophylactic murine model of infection, we found a crucial role of LSD1 in aiding *C. neoformans* mediated modulation of lipid metabolism during pulmonary pathogenesis. Thus, our current investigation sheds light on the novel role of LSD1 in host immunomodulation during *C. neoformans* infection.

## Results

### *C. neoformans* infection promotes aberrant lipid accumulation in macrophages

Formation of FMs involve a combination of the following processes: (i) enhanced uptake of native or modified lipoproteins and lipids, (ii) alteration in intracellular cholesterol and lipid metabolism and (iii) disbalance in homeostasis and transport of lipids ^28^. To begin with, we utilized BODIPY493/503 to visualize total neutral lipid content, and observed elevated levels of LDs in *C. neoformans* infected macrophages with an increase in MOI **(Supplementary Figure 1A, B)** along with an overall increase in the cellular lipid content (**Figure 1 A, B**) and LD formation **(Figure 1C).** In line with our BODIPY 493/503 staining, *C. neoformans* infection resulted in increased expression of genes involved in lipid uptake (*Fat, Msr1, Ldlr*) and lipid droplet biosynthesis (*Plin2*), and no considerable change in the levels of lipid efflux pumps (*Abca1* and *Abcg1*) (**Figure 1D**). Similar results were also observed at protein levels, wherein ADRP (coats lipid droplets), CD36 and LDLR (involved in lipid uptake) were found to be significantly upregulated during *C. neoformans* infection **(Figure 1E)**. Collectively, we observed an active alteration in the host cellular lipid content during *C. neoformans* infection.

**Figure 1:**
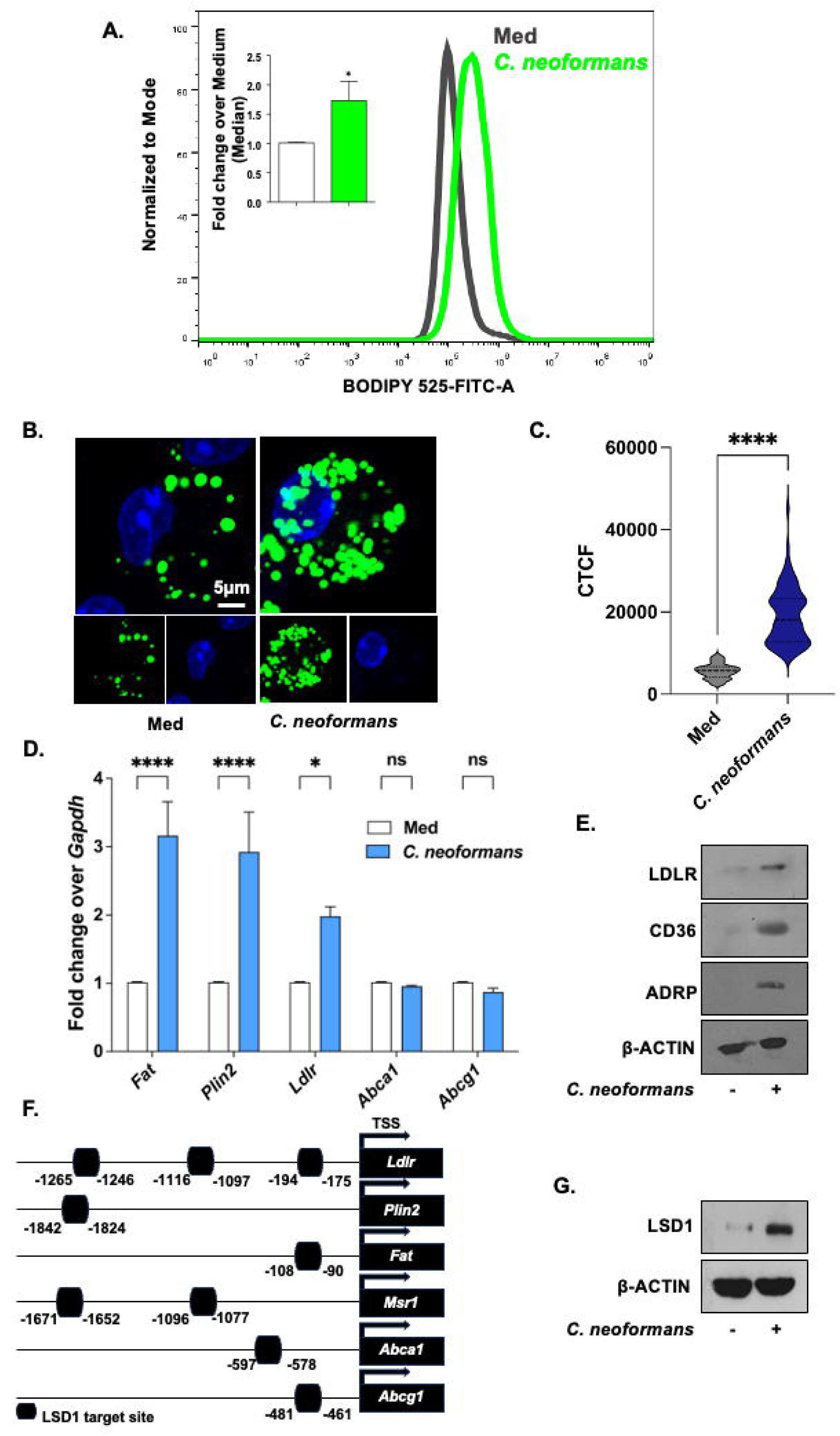
*C. neoformans* infection drives aberrant lipid accumulation in macrophages. **(A-C)** Peritoneal macrophages infected with *C. neoformans* for 48 h at MOI 1:25 and lipid content of the cells was assessed via BODIPY 493/503 staining using **(A)** Flow cytometry and BODIPY 493/503 staining for neutral lipids **(B)** representative image **(C)** quantification. **(D, E)** Peritoneal macrophages infected with *C. neoformans* for 12h at MOI 1:25 and **(D)** transcript levels of indicated genes, **(E)** protein levels of LDLR, CD36 and ADRP by immunoblotting was assessed. **(F)** Promoter analysis for LSD1 Binding site on promoters of indicated genes. **(G)** Peritoneal macrophages were infected with *C. neoformans* for 12 h and the levels of LSD1 by immunoblotting were assessed. MOI: Multiplicity of infection; Med: Medium; CTCF, Corrected Total Cell Fluorescence. ns, non-significant, *, P < 0.05, ****P < 0.0001 (Student’s t-test for A, C and D).

Lipid accumulation arises due to a concerted action of diverse set of genes involved in various pathways that require an intricate transcriptional control ^28^. In this perspective, epigenetic modifiers such as histone modifiers have been implicated to be dynamic under various immune responses and act as important factors in regulating pathogenies and inflammation ^17^. Hence, we were interested to explore the role of epigenetic modifiers in modulating the expression levels of genes involved in foamy macrophages (FM) generation. In this respect, our bioinformatic analysis on the promoters of the genes such *as Ldlr, Acsl, Plin2, Msr1, Fat, Abca1 and Abcg1*, revealed binding sites of diverse transcription factors and epigenetic modifiers. Of interest, LSD1, a well-studied histone demethylase, implicated in regulating lipid metabolism ^22–25^, was found to have a binding site on the promoters of these genes **(Figure 1F).** LSD1, also known as KDMA1, carries out demethylation of H3K9me2 and H3K4me2, resulting in activation or repression of the gene(s), respectively. The dual activity of LSD1 is due to its association with specific transcription factors and co-activators ^29,30^. Examination of protein levels of LSD1 by immunoblotting displayed higher expression in *C. neoformans* infected macrophages as compared to uninfected controls **(Figure 1G).** Therefore, our data suggests a possible role for LSD1 during *C. neoformans* infection.

### Lysine Specific Demethylase-1 (LSD1) fine tunes *C. neoformans*-induced lipid accumulation

We next aimed to assess the contribution of LSD1 on FMs formation during *C. neoformans* infection. To begin with, we analyzed the levels of the genes observed to be implicated in regulating lipid levels for their transcriptional dependency on LSD1. Pharmacological inhibition of LSD1 abolished *C. neoformans*-induced increase in the levels of genes implicated in FM generation, both at transcript and protein levels **(Figure 2 A,B).** Further, upon knocking down LSD1, we observed reduction in the overall lipid content of the *C. neoformans* infected macrophages **(Figure 2C,D)** which was also corroborated upon pharmacological inhibition of LSD1 **(Figure S2 A-C).** In view of the above results, we performed ChIP analysis which revealed an enhanced recruitment of LSD1 over the promoter of genes involved in lipid accumulation, with no significant recruitment over lipid efflux pumps **(Figure 2E**). In line, we also observed a decrease in the levels of H3K9me2 (repressive marks) on the promoters of the genes implicated in FMs formation. Interestingly, unlike the genes associated with lipid accumulation, the levels of repressive H3K9me2 marks remained elevated on the promoters of lipid efflux genes (*Abca1* and *Abcg1*) **(Figure 2E)**. This observation is consistent with our transcript analysis of *Abca1* and *Abcg1*, which showed no significant changes in expression (Figure 1D). These findings suggest that *C. neoformans*-induced LSD1 is selectively recruited to modulate the expression of genes that promote lipid accumulation, thereby facilitating the formation of FMs.

**Figure 2:**
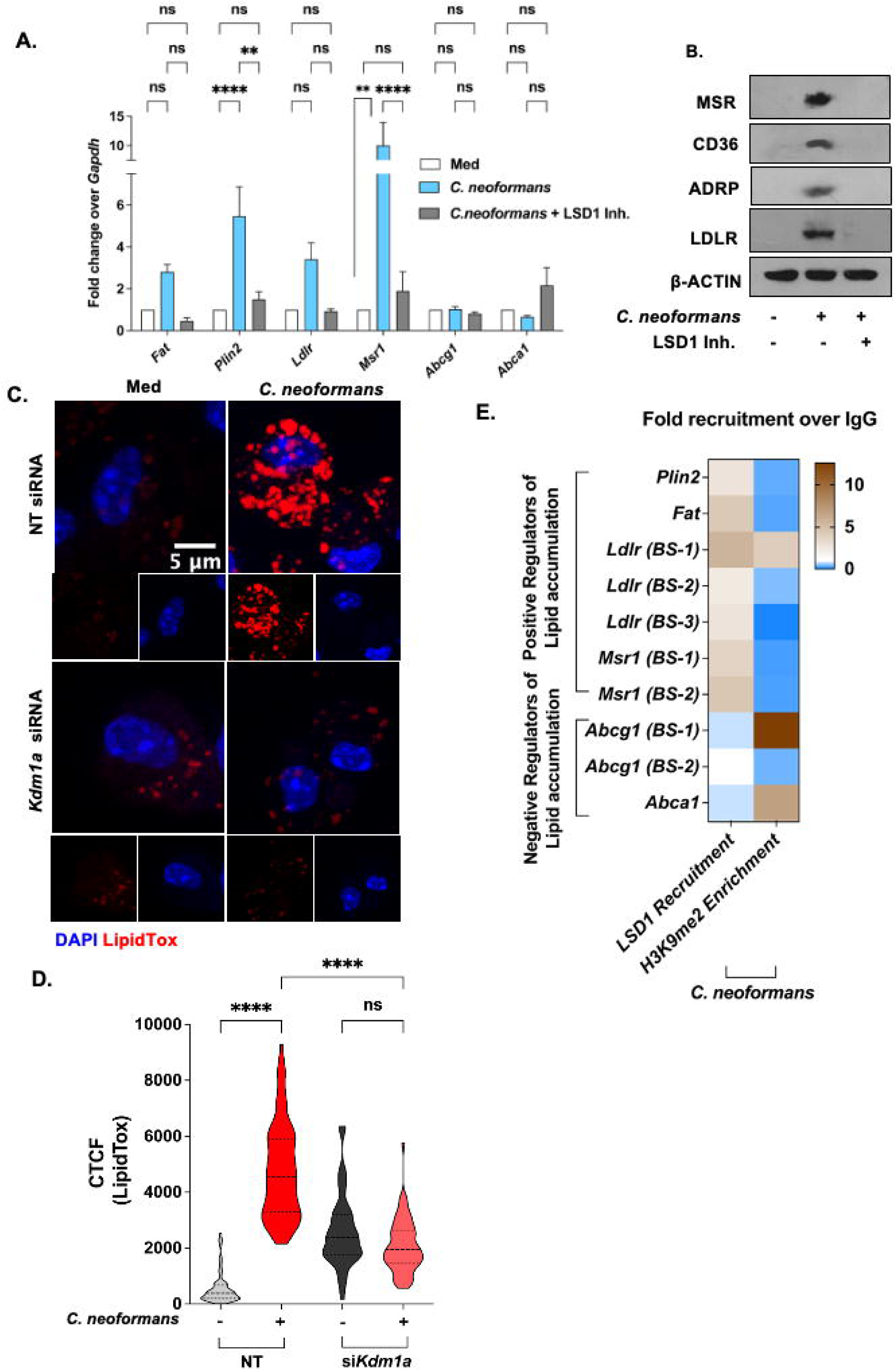
*C. neoformans*-induced LSD1 modulates foam cell formation. **(A, B)** Peritoneal macrophages were infected with *C. neoformans* for 12 h with and without LSD1 Inh (3µM) treatment and indicated genes were assessed at **(A)** transcript levels by qRT-PCR and at **(B)** protein level by immunoblotting. **(C-D)** Mouse peritoneal macrophages transfected with NT or *Kdm1a* siRNA were given 48h *C. neoformans* infection and assessed for lipid droplet accumulation (LipidTOX DeepRed 635/652) **(C)** representative image **(D)** quantification. **(E)** ChIP analysis for the recruitment of LSD1 and enrichment of H3K9me2 over the promoters of indicated genes upon infection with *C. neoformans* for 12 h. MOI for infection is 1:25. Med, Medium; CTCF, Corrected Total Cell Fluorescence. ns, non-significant, **, P < 0.01, ***, P < 0.001, ****, P < 0.0001 (One-way ANOVA for A, D).

### *C. neoformans* induced-LSD1 subdues host’s lipophagy for sustenance of lipid droplets

Autophagy mediated degradation of lipid droplets i.e. lipophagy has been well documented to regulate the total lipid content of the cell. To elucidate the mechanism behind LSD1 mediated lipid dysregulation, we were intrigued to assess whether autophagic degradation of LDs was also getting affected. LSD1 negatively regulates autophagy ^31–35^ and formation of LDs has also been linked to inhibition of autophagy^15^. Autophagy regulates intracellular pool of lipids by employing specific lysosomal degradation of lipid droplets ^36^. Autophagy-mediated lipid degradation has been implicated in various biological context such as adipocyte differentiation, resistance to cell death and regulation of cholesterol efflux ^37^. Moreover, autophagy has been demonstrated to modulate immune response against fungal pathogens, *A. fumigatus* and *C. albicans* ^38,39^. With this premise, we assessed for the probable role of lipophagy in regulating the lipid homeostasis in macrophages upon *C. neoformans* infection. To address this, peritoneal macrophages were infected with *C. neoformans* for 48 h (to permit lipid accumulation), followed by LSD1 inhibition for different time points. Any decrease in lipid content (BODIPY 493/503 staining) post 48 hours upon LSD1 inhibition would indicate active turnover of host lipid content by lipophagy. As shown in **Figure 3 A-B**, LSD1 inhibition for 24h and 18h post 48h of *C. neoformans* infection decreased the augmented lipid levels as observed in *C. neoformans* infected macrophages.

**Figure 3:**
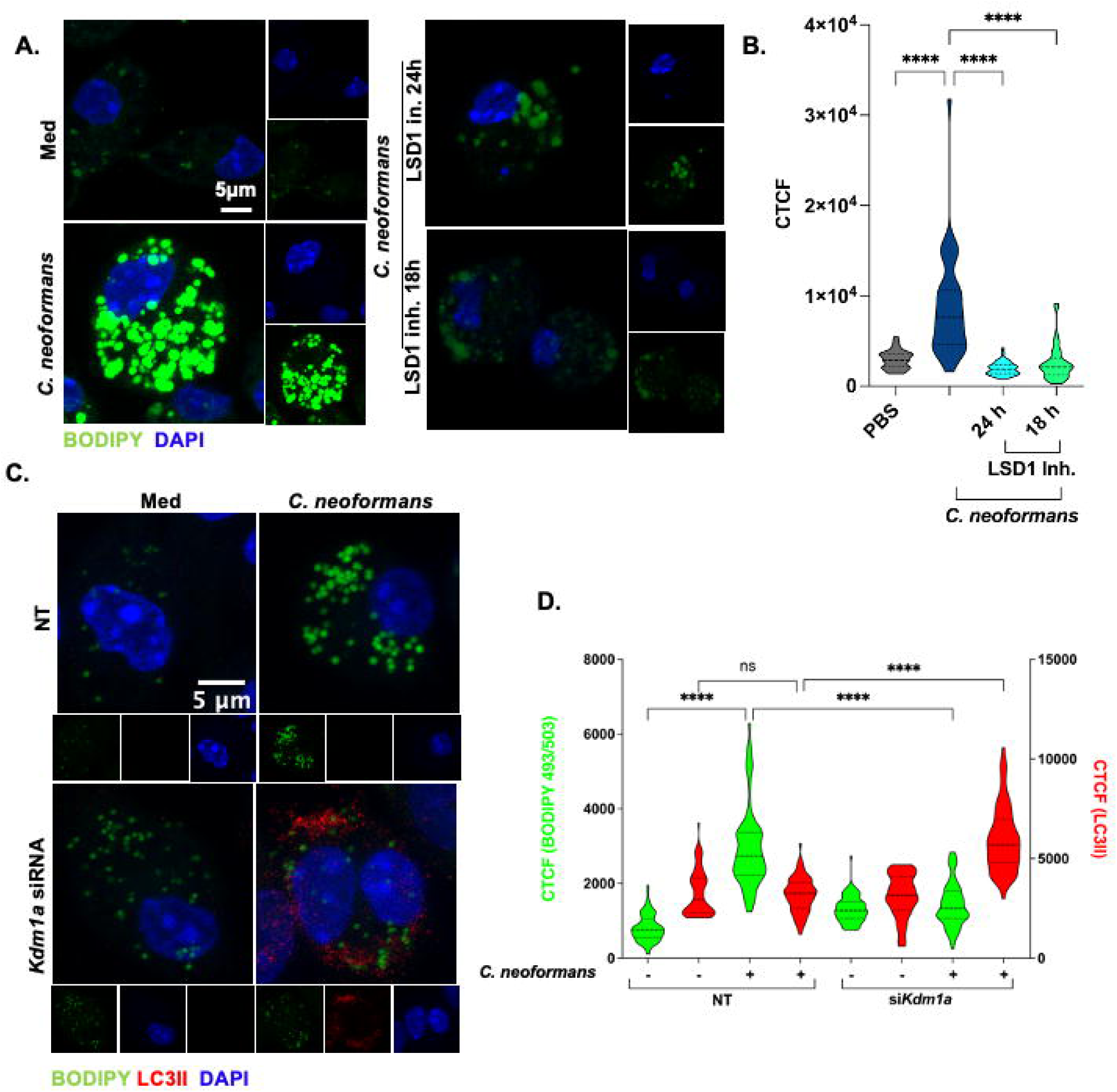
*C. neoformans* induced LSD1 inhibits host’s lipophagy to sustain lipid accumulation. **(A-B)** Peritoneal macrophages were infected with *C. neoformans* for 48 h followed by LSD1 inhibitor (3μM) treatment for indicated time points and stained for neutral lipids by BODIPY 493/503. **(A)** Representative image **(B)** quantification. **(C-D)** Peritoneal macrophages were transfected with NT or *Kdm1a* siRNA and infected with *C. neoformans* for 48 h. Co-staining for neutral lipids (BODIPY 493/503) and LC3II was carried out **(C)** Representative images **(D)** Quantification. MOI for infection is 1:25. Med, Medium; Inh, Inhibitor; CTCF, Corrected Total Cell Fluorescence. ns, non-significant, ****P < 0.0001 (One way ANOVA for B and D).

Further, co-staining for Lipid droplets and autophagy marker (MAP1 LC3B immunostaining) for investigating the possible interplay of the two, yielded an inverse co-relation, implicating autophagy as the cellular process opted for lipid turnover in cells wherein LSD1 was knocked down **(Figure 3 C-D)**or inhibited by pharmacological inhibition of LSD1 **(Figure S3 A, B).** To our surprise, no change was observed in the transcript levels of genes involved in autophagy process **(Figure S3C).** However, in contrast to transcript analysis, protein levels of p62 (autophagy adaptor protein; SQSTM) were compromised in *C. neoformans* infected macrophages wherein LSD1 was inhibited indicating active autophagy **(Figure S3D).**

To further strengthen our hypothesis, we employed mRFP-eGFP-PLIN2 construct ^40^ to assess for the autophagy mediated degradation of lipid droplets. *C. neoformans* infection resulted in the accumulation of mRFP1-EGFP-PLIN2 signal (yellow) highlighting lipid droplets. However, inhibition of LSD1 showed only RFP signal due to quenching of pH sensitive-GFP fluorescence by acidic milieu of lysosomes, confirming an enhanced flux of lipid droplets towards lysosomal mediated degradation **(Figure 4 A, B).** Blocking autophagy by knocking down *Atg5* (involved in autophagy) rescued LSD1 mediated reduction of Lipid droplets consolidating an active role of lipophagy in sustenance of lipid droplets **(Figure 4 C, D; S4A, B).** In conclusion, our results demonstrate that *C. neoformans*-induced LSD1 impedes host lipophagy, thereby sustaining the presence of lipid droplets in macrophages.

**Figure 4:**
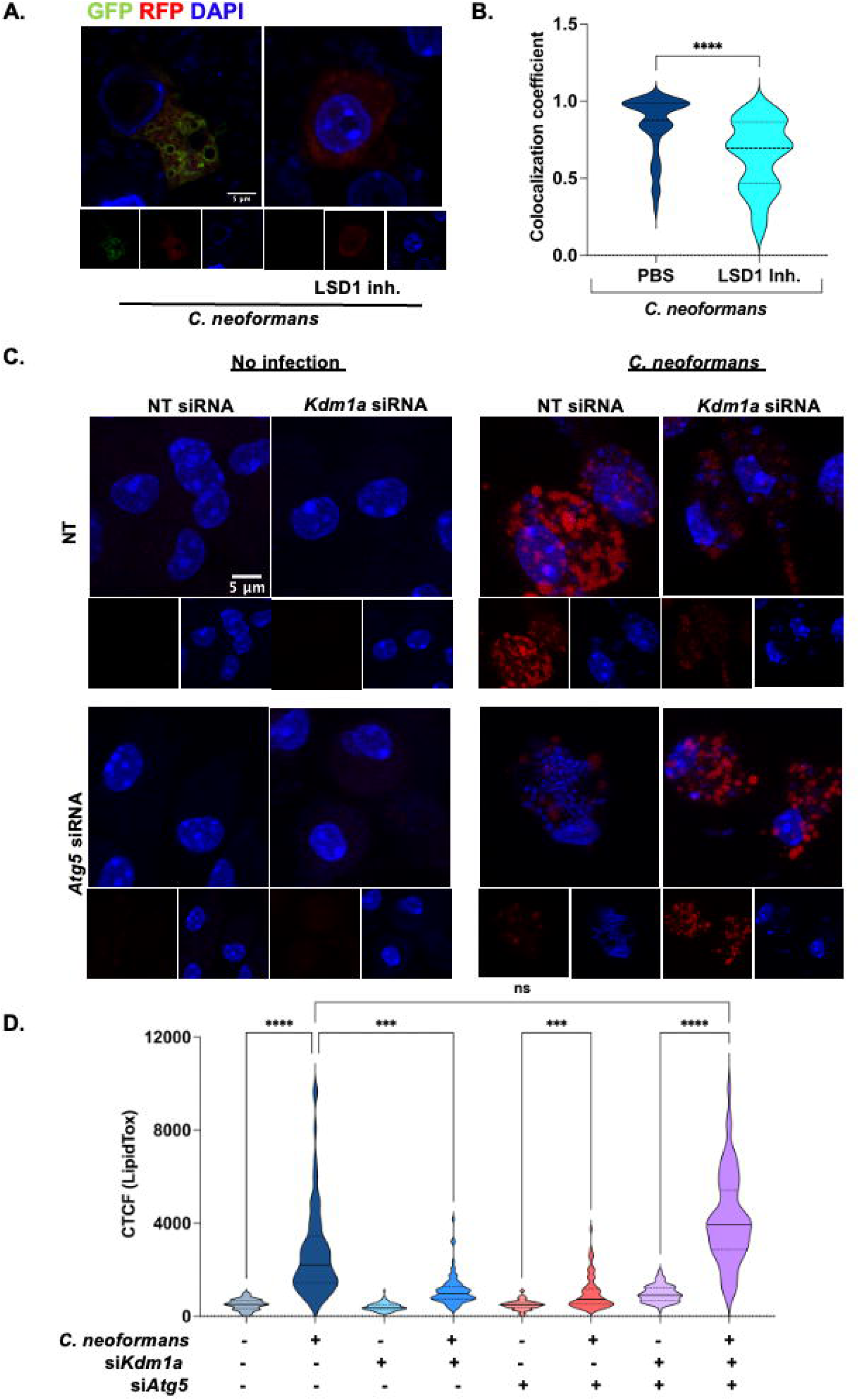
LSD1 inhibition induces lipophagy in *C. neoformans* infected macrophages. **(A-B)** mRFP-eGFP-PLIN2 construct was transfected into RAW264.7 macrophages and infected with *C. neoformans* for 48 h followed by LSD1 inhibitor (3μM) treatment for 18 h. **(A)** Representative images **(B)** Co-localization coefficient. **(C-D)** Peritoneal macrophages were transfected with NT or *Kdm1a* siRNAs in the presence or absence of *Atg5* siRNA followed by infection with *C. neoformans* for 48 h and assessed for lipid droplet accumulation (LipidTOX DeepRed 635/652) **(C)** representative images **(D)** its quantification. NT, non-targeting; Inh, Inhibitor; CTCF, Corrected Total Cell Fluorescence. ****,p<0.0001 (Student’s t-test for B). ns, non-significant ***, p < 0.001 ****,p<0.0001 (One-way ANOVA for D).

### Pyk2-cRaf axis via WNT signaling drives *C. neoformans* induced lipid accumulation

Pathogen Recognition Receptors (PRRs) have an important role in eliciting initial immune responses via activation of signaling pathways in host upon infection ^41^. Hence, we were interested to dissect the molecular signaling at play during *C. neoformans* mediated FMs generation. A role for WNT signaling has been highlighted during *Aspergillus fumigatus* and Candia infections ^42,43^. Moreover, WNT signaling is known to regulate lipid homeostasis via increasing the endosomal flux of Low-Density Lipoproteins (LDLs) ^44^ and is also implicated in driving lipid dysregulation during Mtb pathogenesis ^45^. β-Catenin, a pivotal component of WNT signaling, is regulated by the Ser/Thr kinase GSK3B and β-Catenin is targeted for proteasomal degradation via phosphorylation. Therefore, WNT signaling can be evaluated by an increased inhibitory phosphorylation of GSK3B and a corresponding decrease in phosphorylation levels of β-Catenin. Upon assessing for the same, we found activation of WNT pathway post 1h of *C. neoformans* infection in macrophages **(Figure 5A).**

**Figure 5:**
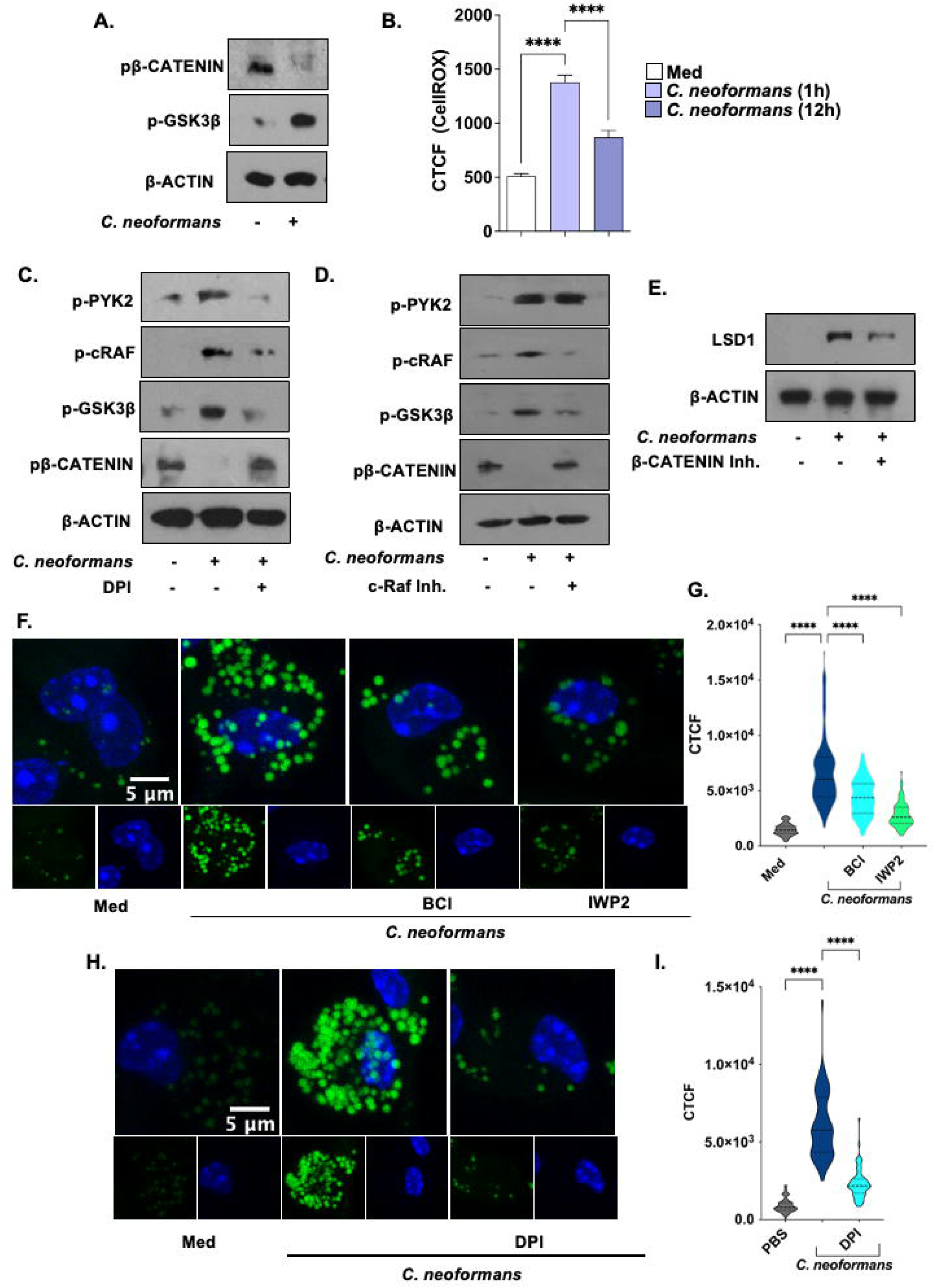
Pyk2-cRaf driven WNT signaling augments lipid content during *C. neoformans* infection. **(A-B)** Peritoneal macrophages infected with *C. neoformans* for 1 h. **(A)** Activation of WNT signaling was checked via immunoblotting. **(B)** CellROX staining for ROS. **(C-D)** Peritoneal macrophages were infected with *C. neoformans* for 1 h in with and without (**C)** DPI or **(D)** cRAF inhibitor treatment and levels of the indicated proteins was assessed by immunoblotting. **(E)** LSD1 levels were assessed upon infection with and without β-Catenin inhibitor treatment (15μM) upon infection with *C. neoformans* for 12 h in peritoneal macrophages. **(F-I)** Peritoneal macrophages infected with *C. neoformans* for 48 h with and without **(F)** β-Catenin inhibitor (15μM) or IWP-II (5mM) or **(H)** DPI treatment and stained with BODIPY 493/503. **(G,I)** quantification. MOI for infection is 1:25. Med, Medium; Inh, Inhibitor; CTCF, Corrected Total Cell Fluorescence. ****P < 0.0001 (One way ANOVA for B, G and I).

Reports suggest increase in ROS (Reactive oxygen species) upon cryptococcal infection ^46^ and can be sensed by tyrosine kinase called Pyk2 ^47,48^. Post 1 hour of *C. neoformans* infection, we found ROS levels to be elevated as seen via CellROX staining **(Figure 5B).** We speculated a role for the ROS-PYK2-cRAF axis in WNT signaling activation as reports evidence cRAF, a serine-threonine kinase, requires activating phosphorylation mark by tyrosine kinase ^49^. Interestingly, *C. neoformans* induced WNT signaling was abrogated upon inhibiting ROS production (via DPI; Diphenyliodenium, ROS inhibitor) **(Figure 5C)** as well as upon inhibition of cRaf **(Figure 5D).**

Further, we found levels of LSD1 to be regulated via WNT signaling as upon β-Catenin inhibition, *C. neoformans* induced expression of LSD1 was abrogated **(Figure 5E).** In line with our previous results, LD accumulation in *C. neoformans* infected macrophages was perturbed upon inhibition of WNT signaling (by β-Catenin inhibitor or IWP-II) **(Figure 5 F, G)** or ROS (by DPI) **(Figure 5 H, I)**. In conclusion, we found ROS-PYK2-cRAF axis to be involved in the activation of WNT signaling and subsequent LSD1 mediated lipid accumulation during *C. neoformans* infection.

### LSD1 aids *C. neoformans* pathogenesis *in vivo*

LSD1 mediated deregulation of host lipophagy intrigued us to investigate its implications during pulmonary pathogenesis of *C. neoformans*. A prophylactic model was utilized wherein mice were administered LSD1 inhibitor intraperitoneally on alternate days followed by intranasal infection with *C. neoformans* **(Figure 6A).** Reduction in the lung fungal burden was observed in mice administered LSD1 inhibitor 4- and 8-days post infection **(Figure 6B)**. In corroboration with the same, alleviation of lung pathology was seen in murine lungs given LSD1 inhibitor treatment **(Figure 6C)** as indicated by granuloma score and percent fraction of lungs having granulomatous lesions **(Figure 6D).**

**Figure 6:**
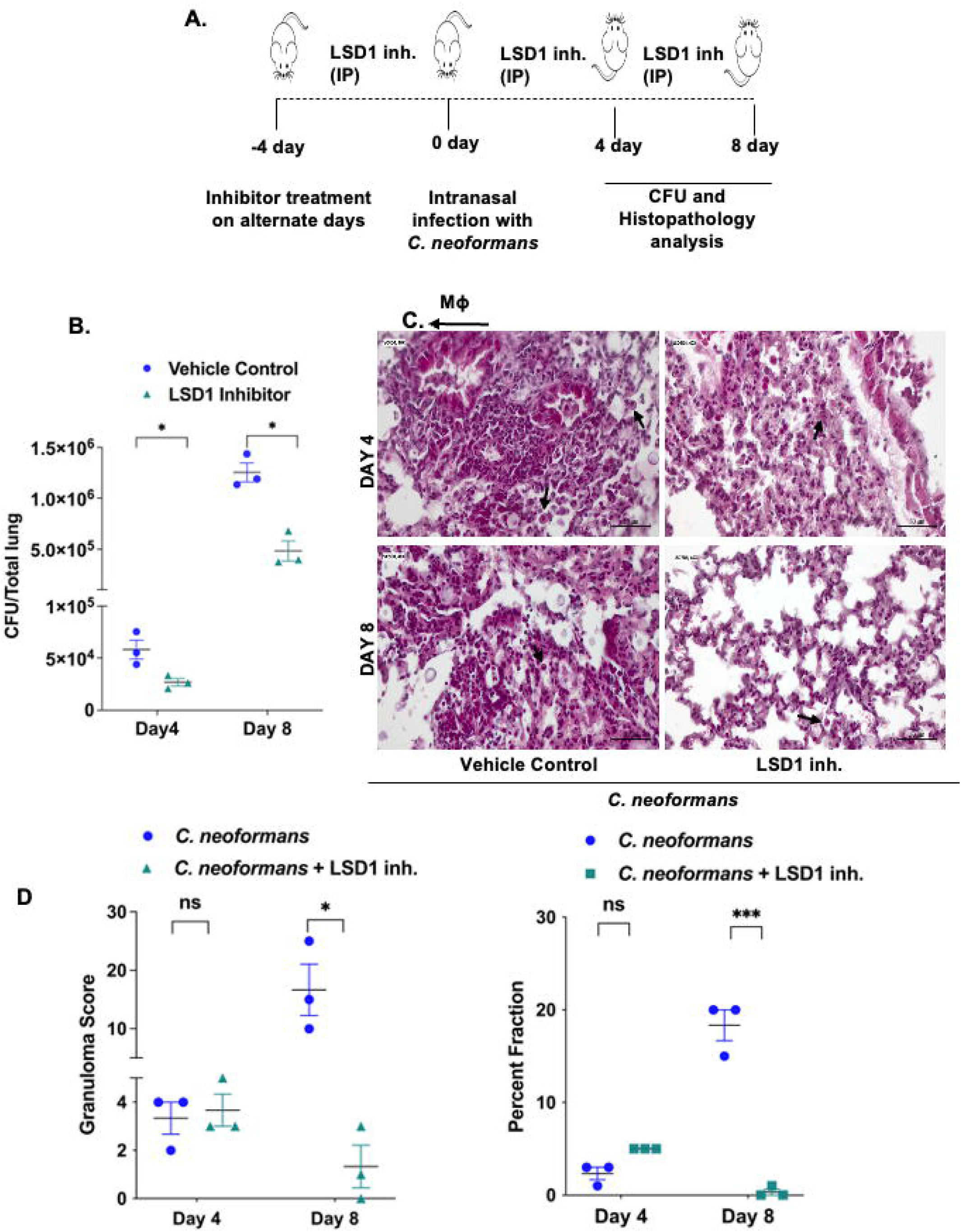
LSD1 supports *C. neoformans* pathogenesis. **(A-D)** Mice were intranasally infected with 10^5^ CFU of *C. neoformans.* **(A)** Schematic representation of *in vivo* mouse model utilized for *C. neoformans* infection with prophylactic treatment of LSD1 inh. (50mg/kg) **(B)** CFU enumeration from the lungs of mice assessed after 4 and 8 days and H&E staining was carried out **(C)** representative image, **(D)** corresponding histological evaluation for granuloma score (left panel) and % fraction of granulomatous area (right panel). N = 2. All data represents the mean ± SEM from 5 mice. inh., inhibitor; ns, non-significant *, P < 0.05 ***, P < 0.001 (Student’s *t*-test for B, D)

Next, we sought out to understand the relevance of LSD1 in lipid dysregulation during pathogenesis **(Figure 7A).** Corroborating our *in vitro* findings, increased expression of most of the genes implicated in FMs formation in the lung of *C. neoformans*-infected mice with a decrease in their levels upon LSD1 inhibition was observed **(Figure 7 B, C).** Examination by immunoblotting of the lung homogenates revealed an upregulation of LDLR, CD36, ADRP in *C. neoformans-*infected murine lungs with a decrease in the levels of ABCA1 which was reversed in lungs of LSD1 inhibitor treated mice **(Figure 7 D, E).**

**Figure 7:**
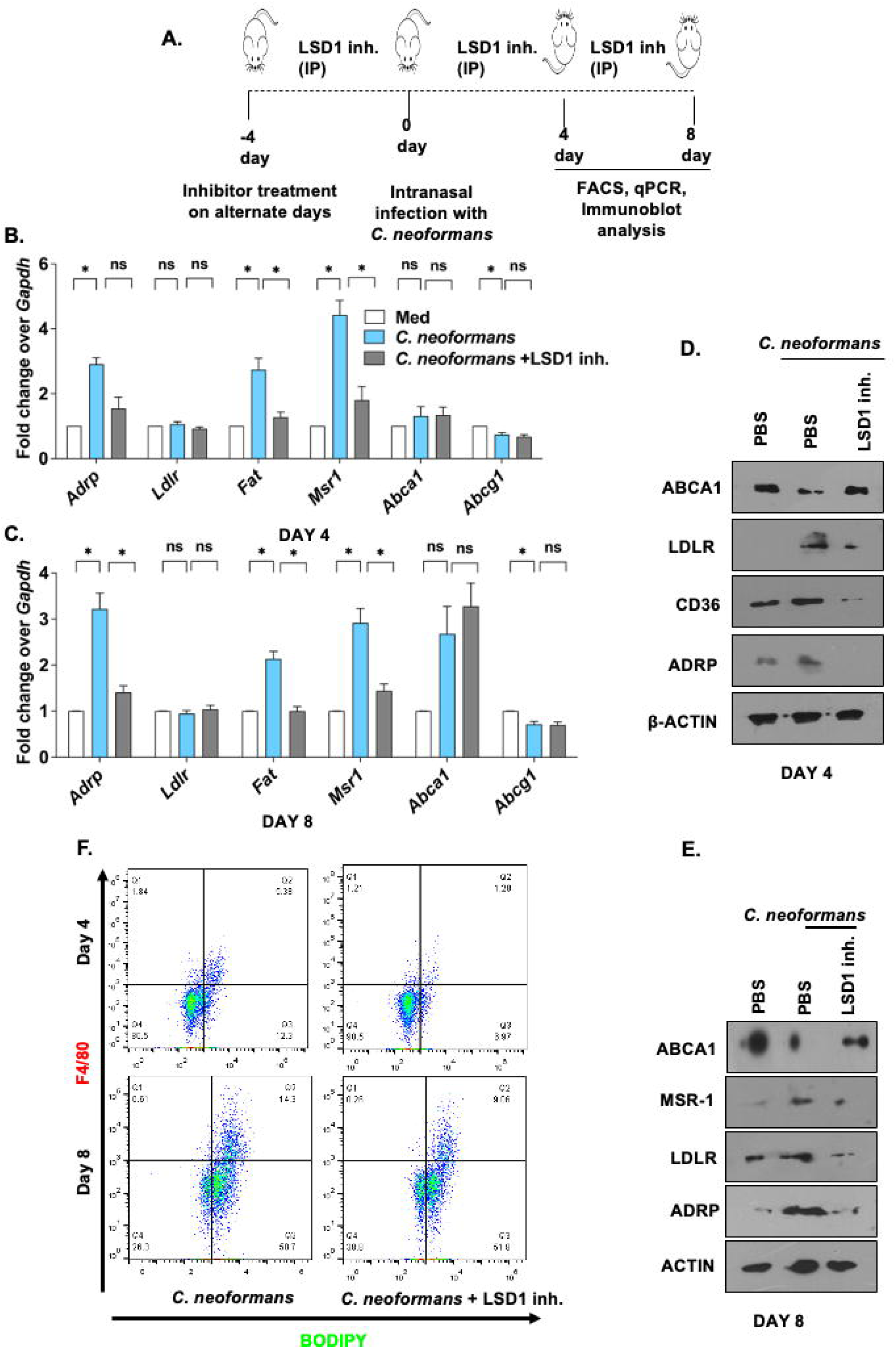
LSD1 regulates lipid accumulation in murine lungs during *C. neoformans* pathogenesis. **(A-F)** Mice were intranasally infected with 10^5^ CFU of *C. neoformans* **(A)** Schematic of *in vivo* mouse model of *C. neoformans* infection with prophylactic treatment of LSD1 inh. The lungs of LSD1 inhibitor (50mg/kg) treated and untreated BALB/c mice after 4 and 8 days of infection were assessed for indicated genes at **(B,C)** transcript levels by qRT-PCR and at **(D,E)** protein level by immunoblotting and **(F)** for lipid content via BODIPY staining by Flow cytometry. inh. inhibitor; ns, non-significant *, P < 0.05 (one way ANOVA for B and C).

Further, we found an increase in the overall lipid content in lung macrophages upon infection, which was decreased in lungs of mice given LSD1 inhibitor treatment at both day 4 and 8 **(Figure 7F)** post *C. neoformans* infection. With these lines of evidence, we highlight a critical role for the epigenetic modifier, LSD1, during pulmonary infection of *Cryptococcus neoformans* via modulating host lipid metabolism.

## Discussion

According to recent reports, 90% of the fungal infections in humans are caused by four species, namely: *Aspergillus, Cryptococcus, Candida*, and *Pneumocystis* ^50^. The fungal priority pathogens list 2022 (WHO FPPL) has categorized *C. neoformans* in the critical group of emerging global health threat of fungal diseases ^51^. Being opportunistic, *C. neoformans* is known to co-exist under various immunocompromised conditions such as in individuals infected with HIV or undergoing immunosuppressive therapies ^52^. Initially manifesting itself as a pulmonary infection, it later spreads to central nervous system wherein it presents itself as cryptococcal meningitis ^53^. Despite such high prevalence of fungal infection stemming from *C. neoformans* infection, molecular insights into its pathogenesis remain elusive.

In the present context, we have shed light on the previously unknown mechanism employed by the *C. neoformans* to induce the formation of lipid-rich FMs and have dissected out the key cellular processes and signalling cascade involved in modulating the lipid accumulation during *C. neoformans* infection. FMs formation involves dysregulation in lipid synthesis and mobilization brought about by different classes of enzymes, receptors and transporters. These foam cells act as a reservoir for the immunomodulatory molecules and site for enzymes such as COX-2 and 5/15-LO. Some of the key players involved in the process include membrane receptors such as Fatty acid translocase (*Fat/*CD36), Low Density lipoprotein receptor (LDLR), Macrophage Scavenging Receptor (*Msr1*) that contribute to the uptake and accumulation of low-density lipoprotein. These lipid molecules-neutral lipids, cholesterol, Triacyl glycerides, once metabolized, can be either sequestered-forming lipid bodies; or effluxed out of the cell *via Abca1* and *Abcg1* (ATP binding cassette transporters) ^6,28^. Thus, upregulation of genes involved in uptake of low-density lipoprotein and down regulation of genes of the lipid efflux pump would be indicative of aberrant lipid accumulation in the cells post infection. Formation of FMs as a mechanism to promote pathogen survival has been well documented in the case of *M. bovis* BCG, *Mycobacterium tuberculosis* and *Chlamydia pneumonia* infection^54,55^. *Encephalitozoon intestinalis* and *Histoplasma capsulatum* are also known to increase production of lipid immunomodulators along with aberrant lipid accumulation in host leukocytes during infection ^56,57^.

In this direction, we have unravelled the role of LSD1, an epigenetic modifier well studied in the context of cancer and reported to modulate immune responses such as the polarization of macrophages, regulating immune checkpoints and restricting viral replication ^58–61^. Herein, we have observed an important role played by LSD1 in fine tuning the expression of genes implicated in lipid uptake and biosynthesis as silencing of LSD1 depleted *C. neoformans*-mediated increase in number of LDs in macrophages. Notably, LSD1 has been highlighted to regulate lipid metabolism in various contexts ^22–25^.

Many reports have shed light on the mechanisms evolved by pathogens to exploit host autophagy machinery in its own favour. Inhibition of autophagy as a successful strategy has been employed by various intracellular pathogens such as *Mycobacterium tuberculosis* ^62^, *Salmonella typhimurium* ^63^ whereas *Shigella flexneri* is known to evade autophagic recognition via masking of its bacterial surface^64^. Some pathogens like *Anaplasma phagocytophilum*^65^, *Yersinia pseudotuberculosis*^66^, *Coxiella burnetii*^67^ induce autophagy but block autophagosome maturation in the host cells to promote its own growth via utilizing the host autophagic vesicle as its own nutrient source^68^. Hepatitis C virus (HCV) have been shown to trigger LD formation and reduction in autophagic catabolism of these LDs via AMPK-SREBP mediated signalling pathway as a mechanism to release viral proteins, infectious viral RNA or HCV virions to escape the host cell^69^.

Several lines of evidence have appended LSD1 in regulating autophagy which can target lipid droplets to lysosomal mediated degradation ^70–72^. Of interest, lipophagy is utilized to maintain the rapid turnover of the lipids in cells^37^. LSD1 was found to inhibit autophagic mediated degradation of lipid droplets, as inhibition of LSD1 augmented lipophagy with concomitant decrease in the total lipid pool of the infected macrophages.

We were further intrigued to understand the initial host immune responses elicited upon infection and signalling axis activated therein to bring about *C. neoformans* mediated foam cell formation. Increase in Reactive Oxygen Species (ROS) as an early immune response has very well been reported ^73^ and is known to regulate various immune signalling pathways during infection^74^, as well as upon *C. neoformans* infection^75^. To our interest, ROS has been reported to modulate WNT signalling pathway, which has previously discussed, plays a pivotal role during fungal infections ^42,76^ and is also involved in modulating lipid droplet homeostasis ^77^. We observed a pivotal role of ROS in activating Pyk2-cRaf resulting in WNT signalling activation. Further investigation showed that inhibition of WNT signalling compromised the levels of LSD1 and consequent lipid accumulation in *C. neoformans* infected macrophages. Further, our *in vivo* infection studies highlight a pro-*C. neoformans* role of LSD1 during pulmonary infection. Host Directed Therapy (HDT) seeks to address crucial host cellular processes altered by pathogens, presenting a promising pathway for developing therapeutic interventions^78^. In this context, various modulatory inhibitors have emerged as potential candidates for drug development. Certain drugs, including EZH2, DOT1L, and BET inhibitors, have shown encouraging outcomes in clinical trials for cancer therapy^79–81^. We propose LSD1 as a potential target for developing HDTs against Cryptococcosis. Inhibition of LSD1 has been posited as a promising strategy against cancer, supported by the development of various LSD1 inhibitor molecules currently undergoing clinical trials ^82–85^.

Altogether, we have implicated the role of WNT driven LSD1 in modulating the lipid profile of the *C. neoformans* infected macrophages by regulating the expression of genes involved in lipid biosynthesis and uptake with simultaneous inhibition of the lipophagy to sustain lipid droplets. We also found out a role of LSD1 in aiding cryptococcal pathogenesis as decrease in the fungal burden and pathology was observed in infected murine lungs upon LSD1 inhibition **(Figure 8).** In this regard, we believe that therapeutic interventions targeting host factors could pave way for prophylactic therapy regime wherein cryptococcosis can be targeted at an early stage itself in immunocompromised individuals.

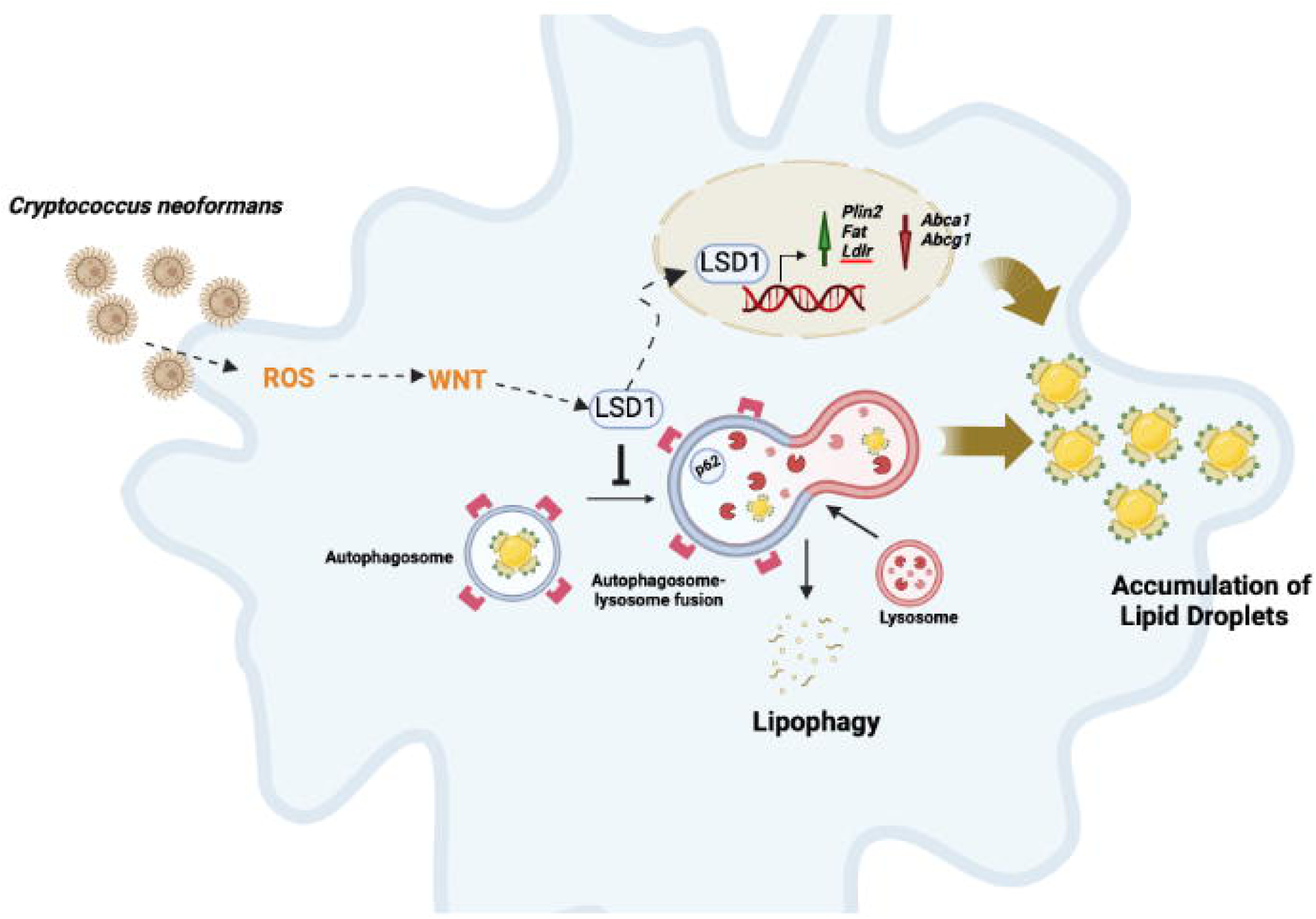

## Materials and Methods

### Ethics statement

Animal experiments (mice) were conducted following approval from the IAEC (Institutional Ethics Committee for Animal Experimentation) and experiments using virulent *Cryptococcus neoformans* (H99-alpha) from the IBSC (Institutional Biosafety Committee). All the protocols for animal care and use were in adherence to the guidelines set by the CCSEA (formerly CPCSEA), (Committee for Control and Supervision of Experiments on Animals), Government of India.

### Mice and cells

BALB/c mice were obtained from Jackson Laboratory and maintained in the Central Animal Facility, Indian Institute of Science (12h light and dark cycle). 4 days after injecting 8% Brewer thioglycolate, macrophages were isolated from the peritoneal exudates of mice. Cells were maintained in DMEM (Dulbecco’s Minimal Eagle Medium) supplemented with 10% heat inactivated Foetal Bovine Serum at 37°C in 5% CO2 incubator. Mouse macrophage cell line, RAW 264.7 was purchased from ATCC for carrying out transient transfection studies using plasmids^86^. PCR based assay was utilized to check the cell lines for mycoplasma.

### Fungi

Wild type strain of *Cryptococcus neoformans var grubii* serotype α was obtained from Prof. Kaustuv Sanyal, Jawaharlal Nehru Centre for Advanced Scientific Research (JNCASR), Bangalore as a kind research gift. The culture was grown in YPD medium (1% yeast extract, 2% Peptone and 2% Dextrose) at 30*°*C overnight. For infection, culture was washed in 1X PBS thrice, counted in hemocytometer and infection was given at indicated MOI.

### *In vivo* infection model

BALB/c mice (n = 40) were subjected to prophylactic administration of LSD1 inhibitor (50mg/Kg) intraperitoneally as mentioned. Mice were anaesthetized using ketamine and xylazine and infected intranasally with *C. neoformans* (CFU 10^5^). Alternately, BALB/c mice were intranasally administered with PBS as control. For assessment, mice were sacrificed at mentioned days, homogenate of the left lobe of the lung was serially diluted in 1X PBS, and plating was done in YPD agar containing to enumerate CFU counts. Other lung lobes were fixed in formaldehyde for further processing.

### Statistical analysis

One-way ANOVA followed by Tukey’s multiple-comparisons degree or Student’s t-test were used for assessing levels of significance. The data represented in the graphs are representative of the values obtained from at least 3 independent experiments. All statistical analyses were carried out using GraphPad Prism 10.0 software (GraphPad Software, USA).

Detailed information concerning the, antibodies, pharmacological reagents, *in vitro* experiments and procedures are given in supplementary.

## Acknowledgments

We thank Central Animal Facility of IISc for providing and maintaining mice for our experimentation. We extend our gratitude to Prof. Kaustuv Sanyal, Jawaharlal Nehru Centre for Advanced Scientific Research (JNCASR), Bangalore for providing *Cryptococcus neoformans* strain and Dr. Mark A. McNiven, Mayo Clinic, Rochester, USA for providing mRFP1-EGFP-PLIN2 construct.

## Funding

This work was supported by funds from the Department of Biotechnology (BT/PR41341/MED/29/1535/2020 DT. 13.08.2021; DBT No.BT/PR27352/BRB/10/163 9/2017, DT.30/8/2018 and BT/PR13522/COE/34/27/2015, DT.22/8/2017 to K.N.B) and the Department of Science and Technology (DST, EMR/2014/000875, DT.4/12/15 to K.N.B.), New Delhi, India. K.N.B. thanks Science and Engineering Research Board (SERB), DST, for the award of J. C. Bose National Fellowship (JBR/2021/000011 and SB/S2/JCB-025/2016). K.N.B. also acknowledges the funding (SP/DSTO-19-0176, DT.06/02/2020) from SERB. The authors thank DST-FIST, UGC Centre for Advanced Study and DBT-IISc Partnership Program (Phase-II at IISc BT/PR27952/INF/22/212/2018), Institute of Eminence (IoE) support of IISc (IE/REDA-23-1757) for the funding and infrastructure support. Fellowships were received from IISc (G.K.L., and A.S.), and Prime Minister’s Research Fellowship (PMRF) (A.S.). The funders had no role in study design, data collection and analysis, decision to publish, or preparation of the manuscript.

## Competing interests

No competing financial interests.

## Abbreviations

FMs: Foamy Macrophages
LDs: Lipid Droplets
LSD1: Lysine Specific demethylase 1
PLIN2: Perilipin 2

## Notes

### Competing Interest Statement

The authors have declared no competing interest.

### Summary of Updates

Figures modified for better representation and summary figure (Figure 8) incorporated

## References

1. Rajasingham R, Govender NP, Jordan A, et al. The global burden of HIV-associated cryptococcal infection in adults in 2020: a modelling analysis. Lancet Infect Dis. 2022;22(12):1748–1755. doi:10.1016/S1473-3099(22)00499-6

2. Bajpai VK, Khan I, Shukla S, et al. Invasive Fungal Infections and Their Epidemiology: Measures in the Clinical Scenario. Biotechnology and Bioprocess Engineering. 2019;24(3):436–444. doi:10.1007/s12257-018-0477-0

3. Casadevall A. Cryptococci at the brain gate: Break and enter or use a Trojan horse? Journal of Clinical Investigation. 2010;120(5):1389–1392. doi:10.1172/JCI42949

4. Freyberg Z, Harvill ET. Pathogen manipulation of host metabolism: A common strategy for immune evasion. PLoS Pathog. 2017;13(12). doi:10.1371/journal.ppat.1006669

5. Van den Bossche J, O’Neill LA, Menon D. Macrophage Immunometabolism: Where Are We (Going)? Trends Immunol. 2017;38(6):395–406. doi:10.1016/J.IT.2017.03.001

6. Guerrini V, Gennaro ML. Foam Cells: One Size Doesn’t Fit All. Trends Immunol. 2019;40(12):1163–1179. doi:10.1016/j.it.2019.10.002

7. Nolan SJ, Fu MS, Coppens I, Casadevall A. Lipids Affect the Cryptococcus neoformans-Macrophage Interaction and Promote Nonlytic Exocytosis. Infect Immun. 2017;85(12). doi:10.1128/IAI.00564-17

8. Bryan AM, Farnoud AM, Mor V, Del Poeta M. Macrophage cholesterol depletion and its effect on the phagocytosis of Cryptococcus neoformans. Journal of Visualized Experiments. 2014;(94). doi:10.3791/52432

9. Hall CJ, Bouhafs L, Dizcfalusy U, Sandstedt K. Cryptococcus neoformans causes lipid peroxidation; therefore it is a potential inducer of atherogenesis. Mycologia. 2010;102(3):546–551. doi:10.3852/08-110

10. Gross NT, Hultenby K, Mengarelli S, Camner P, Jarstrand C. Lipid peroxidation by alveolar macrophages challenged with Cryptococcus neoformans, Candida albicans or Aspergillus fumigatus. Med Mycol. 2000;38(6):443–449. doi:10.1080/MMY.38.6.443.449

11. Guerrini V, Prideaux B, Khan R, et al. Heterogeneity of foam cell biogenesis across diseases. bioRxiv. Published online 2023. doi:10.1101/2023.06.08.542766

12. Olzmann JA, Carvalho P. Dynamics and functions of lipid droplets. Nat Rev Mol Cell Biol. 2019;20(3):137–155. doi:10.1038/s41580-018-0085-z

13. Liu K, Czaja MJ. Regulation of lipid stores and metabolism by lipophagy. Cell Death Differ. 2013;20(1):3–11. doi:10.1038/cdd.2012.63

14. Kounakis K, Chaniotakis M, Markaki M, Tavernarakis N. Emerging roles of lipophagy in health and disease. Front Cell Dev Biol. 2019;7(SEP):467021. doi:10.3389/FCELL.2019.00185/BIBTEX

15. Singh R, Kaushik S, Wang Y, et al. Autophagy regulates lipid metabolism. Nature. 2009;458(7242):1131–1135. doi:10.1038/nature07976

16. Mukherjee T, Bhatt B, Prakhar P, Lohia GK, Rajmani RS, Balaji KN. Epigenetic reader BRD4 supports mycobacterial pathogenesis by co-modulating host lipophagy and angiogenesis. Autophagy. Published online June 28, 2021:1–18. doi:10.1080/15548627.2021.1936355

17. Bayarsaihan D. Epigenetic mechanisms in inflammation. J Dent Res. 2011;90(1):9–17. doi:10.1177/0022034510378683

18. Fol M, Włodarczyk M, Druszczyńska M. Host Epigenetics in Intracellular Pathogen Infections. International Journal of Molecular Sciences *2020, Vol 21, Page* 4573. 2020;21(13):4573. doi:10.3390/IJMS21134573

19. Bierne H, Hamon M, Cossart P. Epigenetics and bacterial infections. Cold Spring Harb Perspect Med. 2012;2(12). doi:10.1101/CSHPERSPECT.A010272

20. Holla S, Prakhar P, Singh V, et al. MUSASHI-Mediated Expression of JMJD3, a H3K27me3 Demethylase, Is Involved in Foamy Macrophage Generation during Mycobacterial Infection. Fortune SM, ed. PLoS Pathog. 2016;12(8):e1005814. doi:10.1371/journal.ppat.1005814

21. Bekkering S, Quintin J, Joosten LAB, Van Der Meer JWM, Netea MG, Riksen NP. Oxidized low-density lipoprotein induces long-term proinflammatory cytokine production and foam cell formation via epigenetic reprogramming of monocytes. Arterioscler Thromb Vasc Biol. 2014;34(8):1731–1738. doi:10.1161/ATVBAHA.114.303887

22. Musri MM, Carmona MC, Hanzu FA, Kaliman P, Gomis R, Párrizas M. Histone demethylase LSD1 regulates adipogenesis. Journal of Biological Chemistry. 2010;285(39):30034–30041. doi:10.1074/jbc.M110.151209

23. Hino S, Sakamoto A, Nagaoka K, et al. FAD-dependent lysine-specific demethylase-1 regulates cellular energy expenditure. Nature Communications *2012 3:1*. 2012;3(1):1–12. doi:10.1038/ncomms1755

24. Li Y, Qian X, Lin Y, et al. Lipidomic profiling reveals lipid regulation by a novel LSD1 inhibitor treatment. Oncol Rep. 2021;46(5). doi:10.3892/OR.2021.8184

25. Abdulla A, Zhang Y, Hsu FN, et al. Regulation of lipogenic gene expression by lysine-specific histone demethylase-1 (LSD1). Journal of Biological Chemistry. 2014;289(43):29937–29947. doi:10.1074/jbc.M114.573659

26. Gu H, Roizman B. Engagement of the Lysine-Specific Demethylase/HDAC1/CoREST/REST Complex by Herpes Simplex Virus 1. J Virol. 2009;83(9):4376–4385. doi:10.1128/jvi.02515-08

27. Maiques-Diaz A, Somervaille TCP. LSD1: Biologic roles and therapeutic targeting. Epigenomics. 2016;8(8):1103–1116. doi:10.2217/epi-2016-0009

28. Witztum JL. You are right too! Journal of Clinical Investigation. 2005;115(8):2072–2075. doi:10.1172/JCI26130

29. Foster CT, Dovey OM, Lezina L, et al. Lysine-specific demethylase 1 regulates the embryonic transcriptome and CoREST stability. Mol Cell Biol. 2010;30(20):4851–4863. doi:10.1128/MCB.00521-10

30. Ding J, Zhang ZM, Xia Y, et al. LSD1-mediated epigenetic modification contributes to proliferation and metastasis of colon cancer. Br J Cancer. 2013;109(4):994–1003. doi:10.1038/BJC.2013.364

31. Ambrosio S, Saccà CD, Amente S, Paladino S, Lania L, Majello B. Lysine-specific demethylase LSD1 regulates autophagy in neuroblastoma through SESN2-dependent pathway. Oncogene. 2017;36(48):6701–6711. doi:10.1038/ONC.2017.267

32. Gu Y, Chen D, Zhou L, et al. Lysine-specific demethylase 1 inhibition enhances autophagy and attenuates early-stage post-spinal cord injury apoptosis. Cell Death Discov. 2021;7(1). doi:10.1038/S41420-021-00455-7

33. He M, Zhang T, Zhu Z, et al. LSD1 contributes to programmed oocyte death by regulating the transcription of autophagy adaptor SQSTM1/p62. Aging Cell. 2020;19(3):e13102. doi:10.1111/ACEL.13102

34. Peixoto P, Grandvallet C, Feugeas JP, Guittaut M, Hervouet E. Epigenetic Control of Autophagy in Cancer Cells: A Key Process for Cancer-Related Phenotypes. Cells *2019, Vol 8, Page* 1656. 2019;8(12):1656. doi:10.3390/CELLS8121656

35. Chao A, Lin CY, Chao AN, et al. Lysine-specific demethylase 1 (LSD1) destabilizes p62 and inhibits autophagy in gynecologic malignancies. Oncotarget. 2017;8(43):74434–74450. doi:10.18632/ONCOTARGET.20158

36. Cuervo AM. Chaperone-mediated autophagy: Dice’s “wild” idea about lysosomal selectivity. Nat Rev Mol Cell Biol. 2011;12(8):535–541. doi:10.1038/NRM3150

37. Liu K, Czaja MJ. Regulation of lipid stores and metabolism by lipophagy. Cell Death Differ. 2013;20(1):3–11. doi:10.1038/CDD.2012.63

38. Tam JM, Mansour MK, Acharya M, et al. The Role of Autophagy-Related Proteins in Candida albicans Infections. Pathogens. 2016;5(2). doi:10.3390/PATHOGENS5020034

39. Martinez J, Malireddi RKS, Lu Q, et al. Molecular characterization of LC3-associated phagocytosis reveals distinct roles for Rubicon, NOX2 and autophagy proteins. Nat Cell Biol. 2015;17(7):893–906. doi:10.1038/NCB3192

40. Schroeder B, Schulze RJ, Weller SG, Sletten AC, Casey CA, McNiven MA. The small GTPase Rab7 as a central regulator of hepatocellular lipophagy. Hepatology. 2015;61(6):1896–1907. doi:10.1002/hep.27667

41. Mogensen TH. Pathogen recognition and inflammatory signaling in innate immune defenses. Clin Microbiol Rev. 2009;22(2):240–273. doi:10.1128/CMR.00046-08

42. Karnam A, Bonam SR, Rambabu N, Wong SSW, Aimanianda V, Bayry J. Wnt-β-Catenin Signaling in Human Dendritic Cells Mediates Regulatory T-Cell Responses to Fungi via the PD-L1 Pathway. mBio. 2021;12(6). doi:10.1128/MBIO.02824-21

43. Zhang Q, Zhang J, Gong M, et al. Transcriptome Analysis of the Gene Expression Profiles Associated with Fungal Keratitis in Mice Based on RNA-Seq. Invest Ophthalmol Vis Sci. 2020;61(6):1–13. doi:10.1167/IOVS.61.6.32

44. Scott CC, Vossio S, Vacca F, et al. Wnt directs the endosomal flux of LDL-derived cholesterol and lipid droplet homeostasis. EMBO Rep. 2015;16(6):741. doi:10.15252/EMBR.201540081

45. Prakhar P, Bhatt B, Lohia GK, et al. G9a and Sirtuin6 epigenetically modulate host cholesterol accumulation to facilitate mycobacterial survival. PLoS Pathog. 2023;19(10):e1011731. doi:10.1371/JOURNAL.PPAT.1011731

46. Coelho C, Bocca AL, Casadevall A. The Intracellular Life of *Cryptococcus neoformans*. Annual Review of Pathology: Mechanisms of Disease. 2014;9(1):219–238. doi:10.1146/annurev-pathol-012513-104653

47. Yang CM, Lee IT, Hsu RC, Chi PL, Hsiao L Der. NADPH oxidase/ROS-dependent PYK2 activation is involved in TNF-α-induced matrix metalloproteinase-9 expression in rat heart-derived H9c2 cells. Toxicol Appl Pharmacol. 2013;272(2):431–442. doi:10.1016/J.TAAP.2013.05.036

48. Lysechko TL, Cheung SMS, Ostergaard HL. Regulation of the Tyrosine Kinase Pyk2 by Calcium Is through Production of Reactive Oxygen Species in Cytotoxic T Lymphocytes. J Biol Chem. 2010;285(41):31174. doi:10.1074/JBC.M110.118265

49. Jelinek T, Dent P, Sturgill TW, Weber MJ. Ras-induced activation of Raf-1 is dependent on tyrosine phosphorylation. Mol Cell Biol. 1996;16(3):1027–1034. doi:10.1128/MCB.16.3.1027

50. Brown GD, Denning DW, Gow NAR, Levitz SM, Netea MG, White TC. Hidden killers: human fungal infections. Sci Transl Med. 2012;4(165). doi:10.1126/SCITRANSLMED.3004404

51. WHO fungal priority pathogens list to guide research, development and public health action. Accessed March 3, 2024. https://www.who.int/publications/i/item/9789240060241

52. Rajasingham R, Smith RM, Park BJ, et al. Global burden of disease of HIV-associated cryptococcal meningitis: an updated analysis. Lancet Infect Dis. 2017;17(8):873–881. doi:10.1016/S1473-3099(17)30243-8

53. Li YN, Wang ZW, Li F, et al. Inhibition of myeloid-derived suppressor cell arginase-1 production enhances T-cell-based immunotherapy against Cryptococcus neoformans infection. Nat Commun. 2022;13(1). doi:10.1038/S41467-022-31723-4

54. Russell DG, Cardona PJ, Kim MJ, Allain S, Altare F. Foamy macrophages and the progression of the human tuberculosis granuloma. Nat Immunol. 2009;10(9):943–948. doi:10.1038/NI.1781

55. Cao F, Castrillo A, Tontonoz P, Re F, Byrne GI. Chlamydia pneumoniae-Induced Macrophage Foam Cell Formation Is Mediated by Toll-Like Receptor 2. Infect Immun. 2007;75(2):753. doi:10.1128/IAI.01386-06

56. Meester I, Rosas-Taraco AG, Solís-Soto JM, Salinas-Carmona MC. The roles of lipid droplets in human infectious disease. Medicina Universitaria. 2011;13(53):207–216. Accessed March 3, 2024. https://www.elsevier.es/en-revista-medicina-universitaria-304-articulo-the-roles-lipid-droplets-in-X1665579611913921

57. Vallochi AL, Teixeira L, Oliveira K da S, Maya-Monteiro CM, Bozza PT. Lipid Droplet, a Key Player in Host-Parasite Interactions. Front Immunol. 2018;9(MAY). doi:10.3389/FIMMU.2018.01022

58. Karakaidos P, Verigos J, Magklara A. LSD1/KDM1A, a Gate-Keeper of Cancer Stemness and a Promising Therapeutic Target. Cancers (Basel). 2019;11(12). doi:10.3390/CANCERS11121821

59. Sobczak M, Strachowska M, Gronkowska K, Karwaciak I, Pułaski Ł, Robaszkiewicz A. Lsd1 facilitates pro-inflammatory polarization of macrophages by repressing catalase. Cells. 2021;10(9). doi:10.3390/CELLS10092465/S1

60. Tan AHY, Tu WJ, McCuaig R, et al. Lysine-Specific Histone Demethylase 1A Regulates Macrophage Polarization and Checkpoint Molecules in the Tumor Microenvironment of Triple-Negative Breast Cancer. Front Immunol. 2019;10(JUN). doi:10.3389/FIMMU.2019.01351

61. Shan J, Zhao B, Shan Z, et al. Histone demethylase LSD1 restricts influenza A virus infection by erasing IFITM3-K88 monomethylation. PLoS Pathog. 2017;13(12). doi:10.1371/JOURNAL.PPAT.1006773

62. Escoll P, Rolando M, Buchrieser C. Modulation of host autophagy during bacterial infection: Sabotaging host munitions for pathogen nutrition. Front Immunol. 2016;7(MAR):81. doi:10.3389/fimmu.2016.00081

63. Tattoli I, Sorbara MT, Vuckovic D, et al. Amino acid starvation induced by invasive bacterial pathogens triggers an innate host defense program. Cell Host Microbe. 2012;11(6):563–575. doi:10.1016/j.chom.2012.04.012

64. Ogawa M, Yoshimori T, Suzuki T, Sagara H, Mizushima N, Sasakawa C. Escape of intracellular Shigella from autophagy. Science (1979). 2005;307(5710):727–731. doi:10.1126/science.1106036

65. Niu H, Xiong Q, Yamamoto A, Hayashi-Nishino M, Rikihisa Y. Autophagosomes induced by a bacterial Beclin 1 binding protein facilitate obligatory intracellular infection. National Acad Sciences. 2012;109(51):20800–20807. doi:10.1073/pnas.1218674109

66. Moreau K, Lacas-Gervais S, Fujita N, et al. Autophagosomes can support Yersinia pseudotuberculosis replication in macrophagesc mi_1456 1108..1123. Wiley Online Library. 2010;12(8):1108–1123. doi:10.1111/j.1462-5822.2010.01456

67. Gutierrez MG, Vázquez CL, Munafó DB, et al. Autophagy induction favours the generation and maturation of the Coxiella-replicative vacuoles. Cell Microbiol. 2005;7(7):981–993. doi:10.1111/j.1462-5822.2005.00527.x

68. Steele S, Brunton J, Kawula T. The role of autophagy in intracellular pathogen nutrient acquisition. Front Cell Infect Microbiol. 2015;5(JUN). doi:10.3389/fcimb.2015.00051

69. Bozza P, D’avila H, Almeida P, Magalhães K, Almeida C, Maya-Monteiro C. Clinical Lipidology Lipid droplets in host-pathogen interactions. Published online 2009. doi:10.2217/clp.09.63

70. Chao A, Lin CY, Chao AN, et al. Lysine-specific demethylase 1 (LSD1) destabilizes p62 and inhibits autophagy in gynecologic malignancies. Oncotarget. 2017;8(43):74434. doi:10.18632/ONCOTARGET.20158

71. Shi Y xu, He Y ji, Zhou Y, et al. LSD1 negatively regulates autophagy in myoblast cells by driving PTEN degradation. Biochem Biophys Res Commun. 2020;522(4):924–930. doi:10.1016/J.BBRC.2019.11.182

72. Ambrosio S, Saccà CD, Amente S, Paladino S, Lania L, Majello B. Lysine-specific demethylase LSD1 regulates autophagy in neuroblastoma through SESN2-dependent pathway. Oncogene *2017 36:48*. 2017;36(48):6701–6711. doi:10.1038/onc.2017.267

73. Kohchi C, Inagawa H, Nishizawa T, Soma GI. ROS and innate immunity. In: Anticancer Research. Vol 29. International Institute of Anticancer Research; 2009:817–822.

74. Li Z, Xu X, Leng X, et al. Roles of reactive oxygen species in cell signaling pathways and immune responses to viral infections. Arch Virol. 2017;162(3):603–610. doi:10.1007/S00705-016-3130-2

75. Coelho C, Souza ACO, Derengowski L da S, et al. Macrophage mitochondrial and stress response to ingestion of Cryptococcus neoformans. J Immunol. 2015;194(5):2345–2357. doi:10.4049/JIMMUNOL.1402350

76. Trinath J, Holla S, Mahadik K, Prakhar P, Singh V, Balaji KN. The WNT Signaling Pathway Contributes to Dectin-1-Dependent Inhibition of Toll-Like Receptor-Induced Inflammatory Signature. Mol Cell Biol. 2014;34(23):4301–4314. doi:10.1128/mcb.00641-14

77. Scott CC, Vossio S, Vacca F, et al. Wnt directs the endosomal flux of LDL-derived cholesterol and lipid droplet homeostasis. EMBO Rep. 2015;16(6):741–752. doi:10.15252/EMBR.201540081

78. Kaufmann SHE, Dorhoi A, Hotchkiss RS, Bartenschlager R. Host-directed therapies for bacterial and viral infections. Nature Reviews Drug Discovery *2017 17:1*. 2017;17(1):35–56. doi:10.1038/nrd.2017.162

79. Groves IJ, Sinclair JH, Wills MR. Bromodomain Inhibitors as Therapeutics for Herpesvirus-Related Disease: All BETs Are Off? Front Cell Infect Microbiol. 2020;10. doi:10.3389/FCIMB.2020.00329

80. Mao Y, Sun Y, Wu Z, et al. Targeting of histone methyltransferase DOT1L plays a dual role in chemosensitization of retinoblastoma cells and enhances the efficacy of chemotherapy. Cell Death & Disease *2021 12:12*. 2021;12(12):1–11. doi:10.1038/s41419-021-04431-y

81. Kim KH, Roberts CWM. Targeting EZH2 in cancer. Nat Med. 2016;22(2):128–134. doi:10.1038/NM.4036

82. Sattler M, Salgia R. LSD1-targeted therapy—a multi-purpose key to unlock immunotherapy in small cell lung cancer. Transl Lung Cancer Res. 2023;12(6):1350–1354. doi:10.21037/TLCR-23-40/COIF

83. Zwergel C, Stazi G, Mai A, Valente S. Trends of LSD1 inhibitors in viral infections. Future Med Chem. 2018;10(10):1133–1135. doi:10.4155/fmc-2018-0065

84. Noce B, Di Bello E, Fioravanti R, Mai A. LSD1 inhibitors for cancer treatment: Focus on multi-target agents and compounds in clinical trials. Front Pharmacol. 2023;14. doi:10.3389/FPHAR.2023.1120911

85. Wass M, Göllner S, Besenbeck B, et al. A proof of concept phase I/II pilot trial of LSD1 inhibition by tranylcypromine combined with ATRA in refractory/relapsed AML patients not eligible for intensive therapy. Leukemia *2020 35:3*. 2020;35(3):701–711. doi:10.1038/s41375-020-0892-z

86. Zhang X, Edwards JP, Mosser DM. The expression of exogenous genes in macrophages: obstacles and opportunities. Methods Mol Biol. 2009;531:123–143. doi:10.1007/978-1-59745-396-7_9

