## Supplemental file for "Histone demethylase LSD1 regulates lipid homeostasis during *Cryptococcus neoformans* infection"

Gaurav Kumar Lohia^a,1^, Awantika Shah^a,1^, Kithiganahalli Narayanaswamy Balaji^a,2^

^a^ Department of Microbiology and Cell Biology, Indian Institute of Science, Bangalore – 560012, Karnataka, India

^1^ Equal contribution

^2^ Corresponding author: Kithiganahalli Narayanaswamy Balaji, Department of Microbiology and Cell Biology, Indian Institute of Science, Bangalore – 560012, Karnataka, India; Ph: +91-80-22933223;

**SUPPLEMENTARY FIGURES AND LEGENDS:**

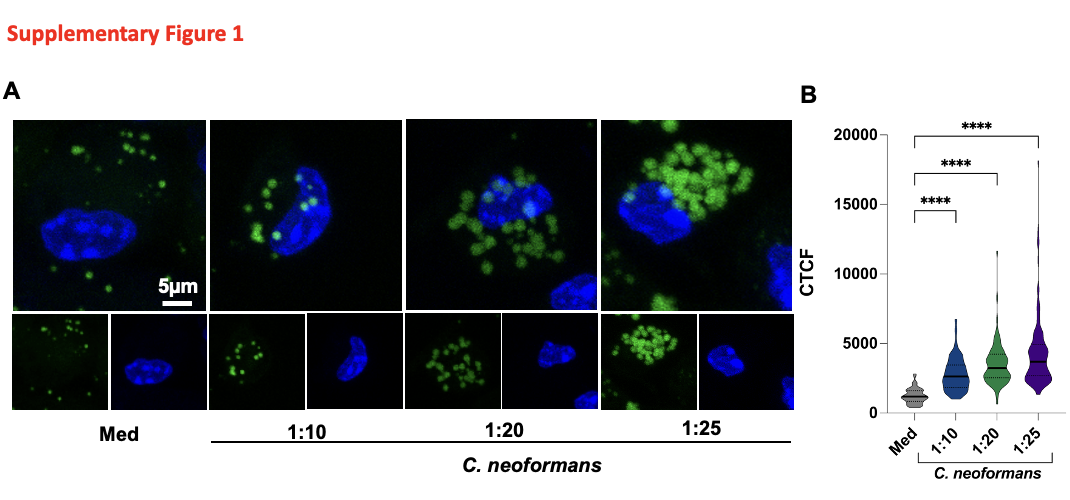

**Supplementary Figure 1: *C. neoformans* infection leads to lipid accumulation in murine macrophages.** Peritoneal macrophages infected with *C. neoformans* for 48 h at the mentioned MOIs and stained with BODIPY 493/503. **(A)** Representative images. **(B)** Quantification. Med, Medium; CTCF, Corrected Total Cell Fluorescence; ****P < 0.0001, (Student’s t-test).

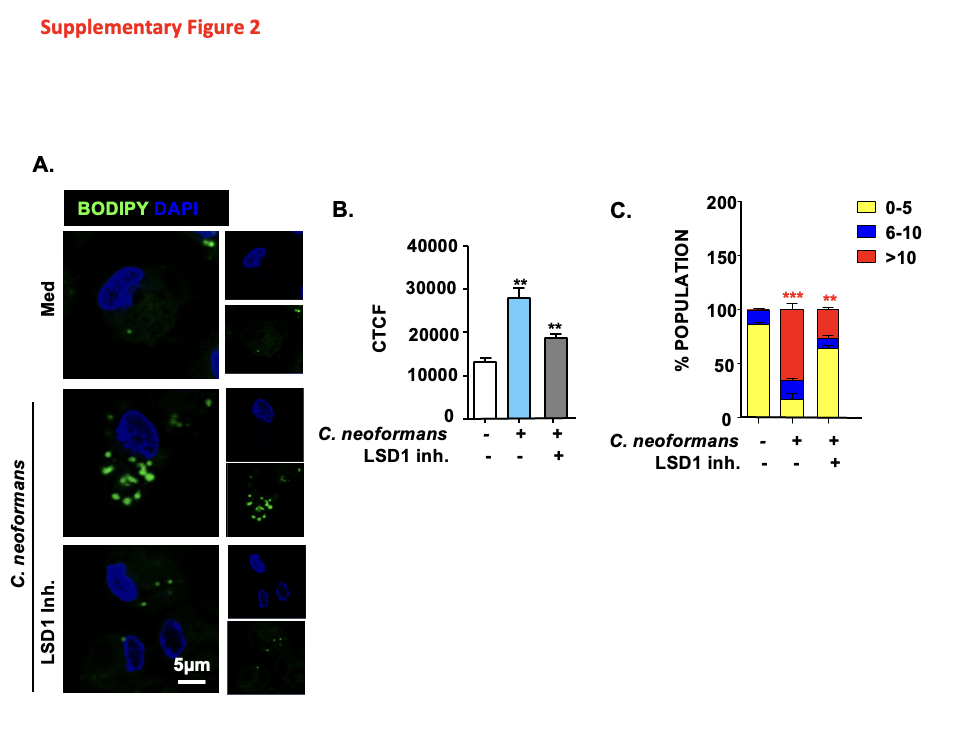

**Supplementary Figure 2: LSD1 regulates foam cell formation during *C. neoformans* infection.** **(A-C)** Peritoneal macrophages were infected with *C. neoformans* for 48 h with and without LSD1 Inh (3µM) treatment and BODIPY 493/503 staining. **(A)** representative images **(B,C)** its quantification. MOI for infection is 1:25. Med, Medium; CTCF, Corrected Total Cell Fluorescence; **, P < 0.01, ***, P < 0.001 (One-way ANOVA for B and C).

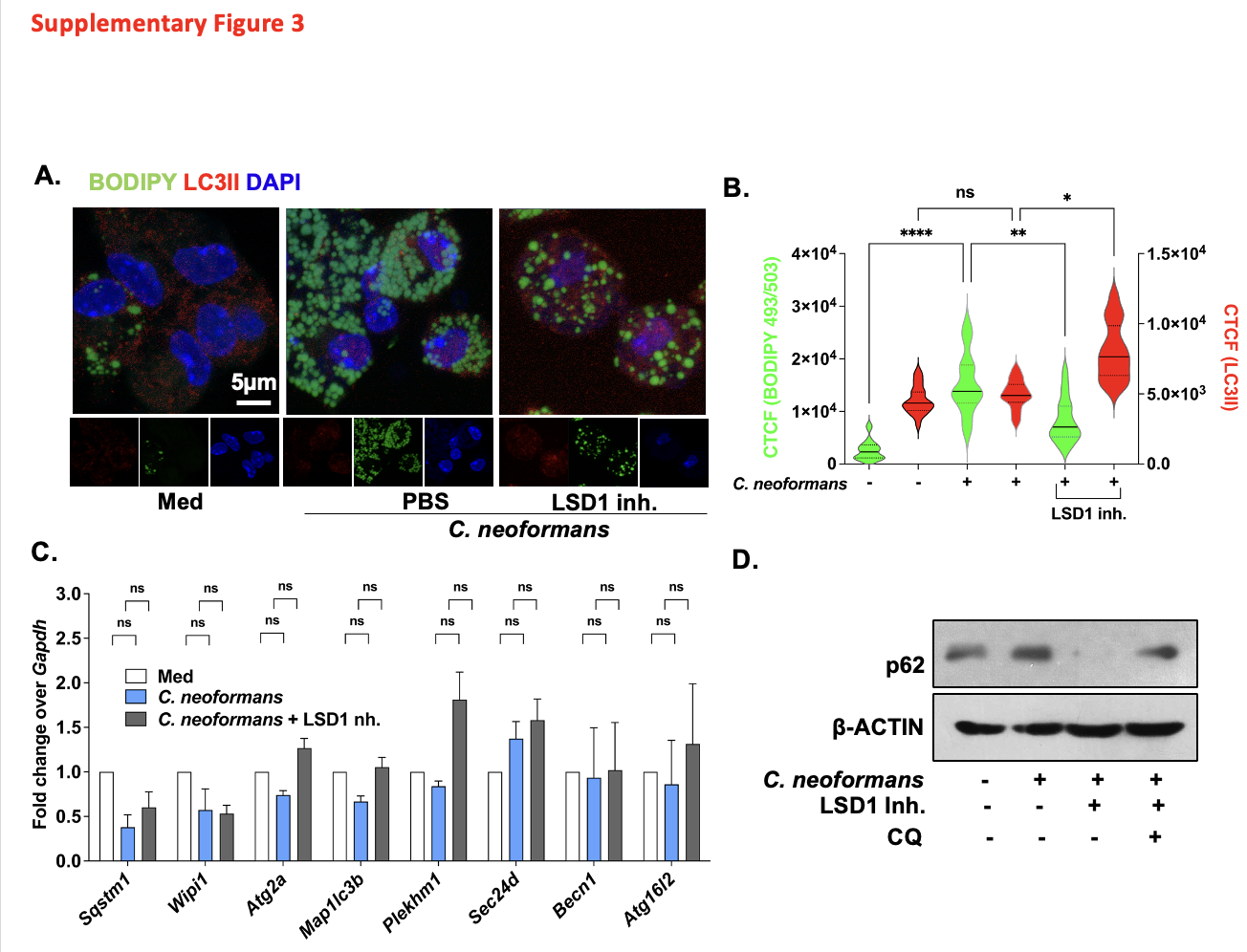

**Supplementary Figure 3: LSD1 inhibits host’s lipophagy to sustain lipid accumulation. (A,B)** Peritoneal macrophages were infected with *C. neoformans* for 48 h followed by LSD1 inhibitor (3μM) treatment for 18h and co-stained for neutral lipids (BODIPY 493/503) and LC3B. **(A)** Representative image **(B)** quantification. **(C)** Peritoneal macrophages were infected with *C. neoformans* in presence and absence of LSD1 inh. for 12 h, **(C)** assessed for the indicated genes by qRT-PCR. **(D)** Peritoneal macrophages were infected with *C. neoformans* for 48 h followed by LSD1 inhibitor (3μM) treatment for 18h in presence and absence of chloroquine (CQ) and assessed for levels of p62 via Immunoblotting. CQ was given 4 h before completion of experiment. MOI for infection is 1:25. Med, Medium; Inh, Inhibitor; CTCF, Corrected Total Cell Fluorescence; ns, non-significant ,*P < 0.05, **P < 0.01, ***P < 0.001, (One way ANOVA for B and C).

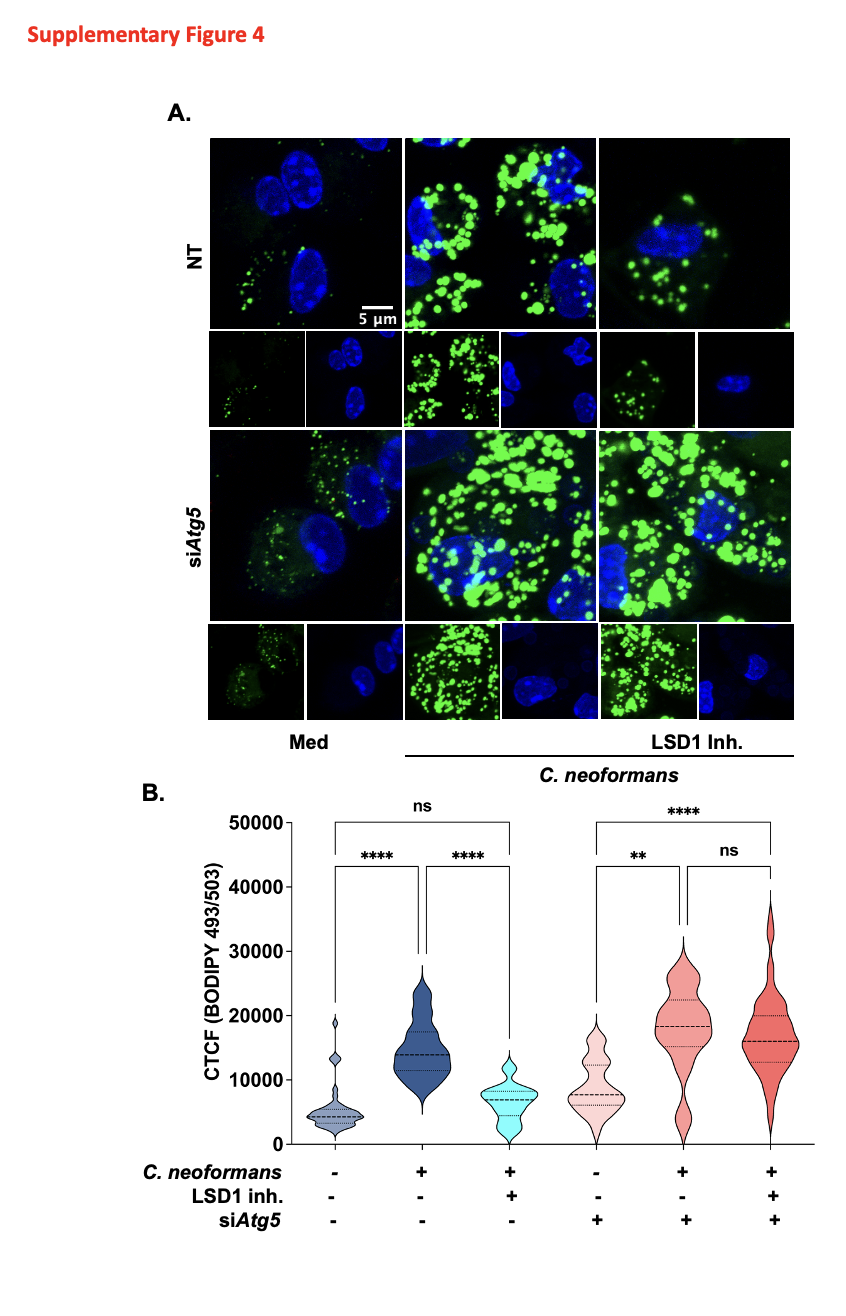

**Supplementary Figure 4: LSD1 inhibits autophagy upon *C. neoformans* infection macrophages to support LD formation. (A,B)** Peritoneal macrophages were transfected with NT or si*Atg5* siRNA using Lipofectamine 3000. Following transfection cells were infected with *C. neoformans* for 48 h followed by LSD1 inhibitor (3μM) treatment for 18 h and assessed for BODIPY **(A)** Representative images **(B)** its quantification. MOI for infection is 1:25. Med, Medium; Inh, Inhibitor; NT, non-targeting; ns, non-significant, **, P < 0.01, ***, P < 0.001 (One way ANOVA for B).

**Materials and Methods**

**Antibodies**

HRP-conjugated anti-β-ACTIN antibody, HRP conjugated anti-rabbit IgG and anti-goat IgG antibody was obtained from Jackson ImmunoResearch (USA). 4′,6-Diamidino-2-phenylindole dihydrochloride (DAPI) from Sigma-Aldrich/ Merck Millipore. Alexa Fluor 660 conjugated CD68 from ThermoFischer Scientific and PE-conjugated F4/80 from Tonbo Biosciences, USA. Cell Signalling and Technology antibodies: anti-H3K9me2, anti-Ser9 phospho-GSK-3β, anti-phospho-β-CATENIN, anti-LSD1, anti-phospho-Pyk2, anti-phospho-c-Raf and anti-p62. Antibodies from Abcam: anti-LDLR, anti-CD36, anti-ADRP, anti-MSR-1, anti-ABCA1. Haematoxylin and Eosin stain solution was from Thomas Baker. BODIPY 493/503 from Molecular Probes (Invitrogen).

**Plasmid**

Dr. Mark A. McNiven, Mayo Clinic, Rochester, USA provided mRFP1-EGFP-PLIN2 construct as a kind research gift.

**Treatment with pharmacological reagents**

Unless otherwise mentioned, for in the *in vitro* experimentations, a pre-treatment of 1h was carried out with concerned pharmacological inhibitors/activators at following concentrations: β-CATENIN inhibitor (15μM), IWP-2 (5μM), LSD1 inhibitor (3µM), DPI (10μM), cRAF inhibitor (2μM), Chloroquine (10 μM) 0.1% DMSO was used as the vehicle control. Viability of the macrophages using the MTT (3-(4,5-Dimethylthiazol-2-yl)-2,5-diphenyltetrazolium bromide) assay was carried out before experimentation.

**Transient transfection studies**

Peritoneal macrophages were transfected with 100nM of either specific siRNAs or non-targeting siRNA (purchased from Dharmacon) and RAW264.7 cells were utilized for carrying out mRFP-eGFP-PLIN2 construct using Lipofectamine-3000 reagent (Thermo Fischer Scientific) in Opti-MEM. Post 36 h (RAW 264.7 cells) or 24h (peritoneal macrophages) of transfection, experiments were carried out as indicated.

**RNA isolation and Real-Time qRT-PCR**

Total RNA was isolated from macrophages using TRI Reagent (Sigma-Aldrich, USA) and 1μg of total RNA was converted into cDNA using First strand cDNA synthesis kit, as per the manufacturer’s protocol. For quantification of target gene expression, quantitative real time PCR was performed with SYBR Green PCR mix (Takara Bio) and the amplification of the gene. *Gapdh* was used as an internal control. To ensure reproducibility of the results, all the reactions were independently repeated at least three times. Ct values were obtained and fold change in transcript levels was calculated using comparative ΔΔCt method. Sequences of primers used in the study are detailed in Supplementary Table 1.

**Immunoblotting analysis**

Cells washed and then scraped with ice-cold 1X PBS were pelleted down via centrifugation. Lysis of the cells was carried out in RIPA buffer [1% NP-40, 0.25% sodium-deoxycholate, 50 mM Tris-HCl pH 7.4, 1 mM EDTA, 150 mM NaCl, 1 mM PMSF, and protease inhibitor cocktail: 1 mM Leupeptin, 1 mM Pepstatin, 1 mM Aprotinin, 1 mM NaF, 1 mM Na3VO4] and incubated for 30min on ice, post which cell lysate was obtained by centrifuging at 13000 rpm for 10 min at 4°C. Protein estimation was carried out using Bradford’s assay and equal concentration of protein lysate for each sample was loaded on SDS-PAGE gel (12% resolving gel, 5% stacking gel) with Tris-Glycine Buffer, pH 8.4. This was followed by transfer onto polyvinylidene difluoride membranes (PVDF) (Millipore) using semi dry western blotting method (Bio-Rad). Blocking to prevent non-specific binding was carried out using 5% non-fat dry milk powder in TBST [20 mM Tris-HCl (pH 7.4), 137 mM NaCl, and 0.1% Tween 20] for 1h and incubated with primary antibody (diluted in 5% BSA) at 4˚C overnight. Membrane was then blocked with to 5% non-fat milk in TBST for 1h. Following blocking, blots were washed in TBST and then incubated with goat anti-rabbit IgG secondary Ab conjugated to HRP (Jackson Immunologicals, USA) (diluted in 5% non-fat milk made in TBST). Immunoblots were developed using enhanced chemiluminescence protocol (Bio-Rad) with β-ACTIN as loading control.

**Chromatin Immunoprecipitation (ChIP) Assay**

Post infection, to crosslink the DNA-Protein interactions of the cells, macrophages were fixed with 3.7% formaldehyde (15min, RT) followed by addition of 125 mM glycine to quench the reaction. Lysis of the nuclei was carried out in 0.1% SDS lysis buffer [50 mM Tris-HCl (pH8.0), 10 mM HEPES (pH 6.5), 200 mM NaCl, 10mM EDTA, 0.5 mM EGTA, 0.1% SDS, 1 mM PMSF, 1 μg/ml of each pepstatin, aprotinin, leupeptin, , 1 mM Na3VO4 and 1 mM NaF] followed by shearing of chromatin at high power of 70 rounds: 30 sec pulse ON/45 sec OFF using Bioruptor Plus (Diagenode). Specific antibody or anti-rabbit IgG complexed with Protein A agarose beads (Bangalore Genei) was used to immunoprecipitated DNA fragments of an average size of 500 bp. Next, sequential washing steps of the immunoprecipitated complexes were carried out thrice in each buffer [Wash Buffer A: 50 mM Tris-HCl (pH8.0), 1 mM EDTA, 500 mM NaCl, 0.1% Sodium deoxycholate, 1% Triton X-100, 0.1% SDS and protease inhibitors; Wash Buffer B: 50 mM Tris-HCl (pH 8.0), 250mM LiCl, 0.5% Sodium deoxycholate, 1 mM EDTA, 0.5% NP-40, and protease inhibitors; TE buffer: 10 mM Tris-HCl (pH 8.0), 1 mM EDTA] followed by elution in buffer: 0.1 M NaHCO3, 1% SDS. Degradation of RNA and Protein was done with RNase A and Proteinase K. Phenol-chloroform-ethanol extraction method was used for DNA precipitation. Quantitative real time PCR analysis was carried out with all values in the test samples normalized to amplification of the specific gene in Input and IgG pull down. Graphs were represented as fold change in modification or enrichment. Primer sequences used for ChIP assay are given in Supplementary Table 2.

**BODIPY 493/503 lipid staining**

Cells on the coverslips were fixed with 3.7% formaldehyde for 15min followed by three washes with 1X PBS. Cells were stained with 10μg/ml BODIPY493/503 for 30 min in dark. After 1X PBS washes, nucleus staining with DAPI for 5 min and mounted on a slide with glycerol. Confocal images were captured with Zeiss LSM 710 Meta confocal laser scanning microscope (Carl Zeiss AG, Germany) using a plan-Apochromat 63X/1.4 Oil DIC objective and analyzed using ZEN 2009 software. The lipid bodies were counted in over 100 cells from different fields. Frequency of populations with different number of lipid bodies (0-5 in yellow, 6-10 in blue, >10 in red) were plotted in terms of percentage. Free hand selection tool in ImageJ was used to select cells and measure the area-integrated intensity and mean grey value of the Z-stacks. The area without fluorescence was used to calculate the background values. the following formula was used to calculate Corrected Total Cell Fluorescence (CTCF) = Integrated intensity – (area of selected cell X Mean florescence of background reading).

**Immunofluorescence (IF) along with BODIPY staining**

Post fixation with 3.7% formaldehyde and washes with 1X PBS, non-specific binding was blocked via incubating the cells with 2% BSA containing 0.02% saponin in PBST for 1h. Following this, cells were stained with specific primary antibody (as mentioned in the text) at 4ºC overnight. Cells were then stained with Alexa647-conjugated secondary antibody and 10μg/ml BODIPY493/503 in dark for 1h at room temperature and nuclei were stained with DAPI and mounted with glycerol. Confocal imaging and analysis was carried out as described above.

**CellROX Oxidative Stress Reagent staining**

CellROX Deep Red Reagent obtained from Thermo Fisher Scientific, USA was used as per manufacturer’s instructions. Briefly, CellROX Deep Red Reagent at a final concentration of 5 mM in DMEM without serum was used to stain peritoneal macrophages at 37 °C for 30 min. Cells were washed with 1X PBS thrice and fixed for 15min with 3.6 % formaldehyde. Post, nuclei staining with DAPI, images were captured and analysed as described above.

**BODIPY staining via Flow Cytometry**

Peritoneal macrophages were gently scraped in 1X PBS or single cell suspension from lung homogenate was prepared. Briefly, Murine lungs were harvested and washed in 1X PBS and minced using scissors (6ml RPMI -/-) subjected to Collagenase and DNASe-I treatment (37°C, 75min). 3ml heat Inactivated FBS was added and suspension passed through cell strainer (70 micron) and Centrifuged at 300g for 5min, 10°C. To the cell pellet, RBC lysis buffer was added (2min,3ml) and centrifuged at 300g for 5min, 10°C. Post washes with 1X PBS wash, cells resuspended in FACs buffer (5ml) (FBS-HI, 0.5M EDTA, 1XPBS) and cell suspension prepared. Post this, live/dead staining was carried out via using Propidium Iodide (PI) at a concentration of 5μg/ml for 5min at room temperature. Post washes with 1X PBS, cells were stained with macrophage marker F4/80 for 45min at 4°C. Cells were fixed with 3.6% formaldehyde for 15 min at room temperature followed by staining with 10μg/ml BODIPY493/503 for 30 min in dark. Cells were subjected to flow cytometry analysis using BC cytoFLEX and analysed using FlowJo software.

**Hematoxylin and Eosin (H&E) staining**

Using Leica RM2245 microtome, 5 μm thick sections were obtained from formalin-fixed, paraffin-embedded mouse lung tissue samples. Deparaffinization and rehydration of the sections was carried out followed by Hematoxylin and Eosin staining. The sections were then dehydrated and mounted with coverslip using permount. A blinded analyses was carried out by a pathologist to obtain granuloma scores and fraction of lung involved in pathology.

**Supplementary Tables:**


**Supplementary Table 1: List of primers for mouse gene expression analyses**

| ***Genes*** | **Forward primer (5’-3’)** | **Reverse primer (5’-3’)** |
| --- | --- | --- |
| *Gapdh* | gagccaaacgggtcatcatct | gaggggccatccacagtctt |
| *Abca1* | aaaaccgcagacatccttcag | cataccgaaactcgttcaccc |
| *Abcg1* | gtggatgaggttgagacagacc | cctcgggtacagagtaggaaag |
| *Plin2* | ggagtggaagagaagcatcg | tggcatgtagtctggagctg |
| *Fat* | cagtcggagacatgcttattgag | tttgccacgtcatctgggttt |
| *Ldlr* | tgactcagacgaacaaggctg | atctaggcaatctcggtctcc |
| *Msr1* | ttcactggatgcaatctccaag | ctggacttctgctgatactttgt |
| *Sqstm1* | aggatggggacttggttgc | tcacagatcacattggggtgc |
| *Becn1* | aggcgaaaccaggagagac | cctccccgatcagagtgaa |
| *Map1lc3b* | ttatagagcgatacaagggggag | cgccgtctgattatcttgatgag |
| *Atg2a* | gtgtggtactacgggaggtct | cctggtgttgccgtccaat |
| *Plekhm1* | cagaaacagcatgtgtctctgg | ggtgagtgtgcttcttcctttt |
| *Atg16l2* | ggagagactcagtccaaggaa | ccacgtcattgcagtaggaaag |
| *Sec24d* | agcctggaatgggtatctctc | gtgccacactattgacaggtg |
| *Wipi1* | ctgcttctctttcaaccaagact | acgtcagggatttcattgctt |

**Supplementary Table 2: List of primers for ChIP Assays**

| **Gene Name** |  | **Sequence** |
| --- | --- | --- |
| **For LSD1 binding (5’-3’)** | | |
| *Plin2* | Forward | atctttgcaatgcagctttc |
|  | Reverse | aacctggtgccacaatatc |
| *Ldlr (1)* | Forward | gtctttgacaggcagatgga |
|  | Reverse | tgtgagttgatttctctcggt |
| *Ldlr (2)* | Forward | agttagaggcaggcagat |
|  | Reverse | cctcctcttctttccttctttc |
| *Ldlr (3)* | Forward | agaggaggaggaaacagaat |
|  | Reverse | ggtaggacccagtagttagaa |
| *Msr1 (1)* | Forward | cttctactgggttgccttg |
|  | Reverse | gcttcacattacctctactctc |
| *Msr1 (2)* | Forward | cctcctttagtccacatggt |
|  | Reverse | cacacacacacactgtcttta |
| *Abca1* | Forward | tggttgtaaagcagcaagt |
|  | Reverse | tatgcagcttccctcatttc |
| *Abcg1 (1)* | Forward | caaaggaatgtgaatcaaaggg |
|  | Reverse | gtatcagcatgcctggttt |
| *Abcg1 (2)* | Forward | gtgatgctagggagaagcttag |
|  | Reverse | ctgttgaaatgattctctgtcacc |
| *Fat* | Forward | acactcatattccatggtgaag |
|  | Reverse | ttgaacttccaagtgctgag |
